# Behavioral and neural alterations of the ventral tegmental area by exposure to junk food in rats

**DOI:** 10.1101/2024.10.14.618156

**Authors:** Jaume F. Lalanza, John C. Oyem, Patty T. Huijgens, James E. McCutcheon, Roy Heijkoop, Eelke MS Snoeren

## Abstract

The brain reward system is essential for regulating appetitive and consummatory behaviors in response to various incentive stimuli. Junk food, characterized by its high palatability, is particularly associated with the potential for excessive consumption. While prior studies indicate that excessive junk food intake can impact reward circuitry, the precise mechanisms underlying these effects remain elusive. Furthermore, it is unclear whether the functionality of this neural system is similarly altered in response to other natural rewards. In this study, we used fiber photometry combined with a behavioral reward test to investigate the effects of six weeks of excessive cafeteria (CAF) diet consumption on ventral tegmental area (VTA) neural activity and behavioral responses to food and sexual rewards in female rats. Our findings demonstrate that prolonged exposure to a CAF diet reduced the exploration and consumption of food rewards. These behavioral changes were accompanied by attenuated neural activity in the VTA. Similarly, reductions in VTA activity were observed in response to a sexual partner, although no significant behavioral differences were detected during sexual interactions. Moreover, a two-week reversal diet of standard chow proved insufficient to restore VTA neural activity in CAF-exposed animals, which continued to exhibit decreased VTA responses to both food rewards and sexual partners. Our results suggest that prolonged junk food exposure induces desensitization of the VTA, resulting in reduced responsiveness to natural rewards.

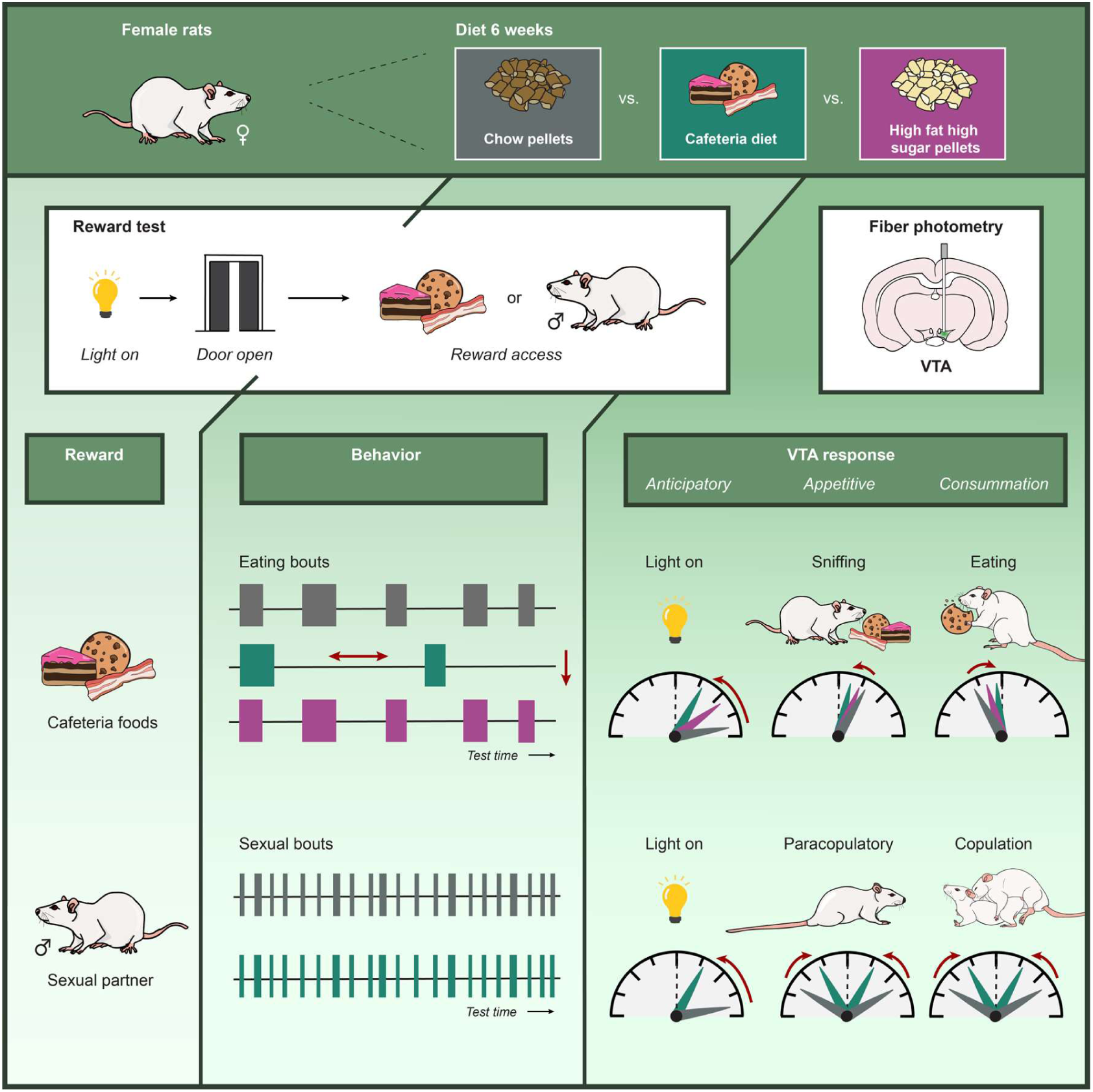

## 1. Introduction

The widespread availability and excessive consumption of processed foods rich in energy, salt, fat and sugar, also called junk food, is linked to the increased prevalence of obesity and overweight globally (Dicken & Batterham, 2024; Malik, Willett, & Hu, 2013; Morris, Beilharz, Maniam, Reichelt, & Westbrook, 2015; WHO, 2024). The highly palatable characteristics of these foods activate hedonic and motivational mechanisms, promoting food-seeking behavior and overeating (Morales & Berridge, 2020; Morris, et al., 2015). Modern energy-rich diets are consumed not only for nutritional content but also for their hedonically pleasant traits (Berridge, Ho, Richard, & DiFeliceantonio, 2010; Ulug, Pinar, & Yildiz, 2025; Zheng & Berthoud, 2007). Besides the risk for obesity, junk food consumption is also associated with a high risk for other diseases such as cardiovascular disease, type 2 diabetes, infertility, hypertension and some cancers (Briggs, Petersen, & Kris-Etherton, 2017; Elias, Elias, Sullivan, Wolf, & D’Agostino, 2005; Guh, et al., 2009; Lane, et al., 2024; Leggio, et al., 2017; Oikonomou, et al., 2018), and cognitive and neurological changes are also found to be linked to junk food consumption (Davidson, et al., 2013; Noble, Hsu, & Kanoski, 2017; O’Brien, Hinder, Callaghan, & Feldman, 2017; Raji, et al., 2010; Tsan, Decarie-Spain, Noble, & Kanoski, 2021).

Excessive junk food consumption can also disrupt neural reward mechanisms (Garcia-Garcia, et al., 2014; Nummenmaa, et al., 2012; Stuber, Schwitzgebel, & Luscher, 2025). Under normal circumstances, dopamine transmission from the ventral tegmental area (VTA) to the nucleus accumbens (NAc) partly regulates food palatability and consumption (Berridge & Robinson, 2003; Yoshida, et al., 1992). As such, VTA dopamine neurons regulate many aspects of reward-related behavior including reinforcement, decision making, attention, and impulsivity (Fields, Hjelmstad, Margolis, & Nicola, 2007; Flores-Dourojeanni, et al., 2021; Morales & Margolis, 2017; Solie, Girard, Righetti, Tapparel, & Bellone, 2022). They predict reward value and integrate past experiences with effort required to obtain rewards (Berke, 2018; D’Ardenne, McClure, Nystrom, & Cohen, 2008; Eshel, Tian, Bukwich, & Uchida, 2016; Hamid, et al., 2016; Schultz, 2007, 2016; Syed, et al., 2016). Acute exposure to palatable foods increases dopamine release in the NAc (Bassareo & Di Chiara, 1999; Hajnal, Smith, & Norgren, 2004; Martel & Fantino, 1996a; Roitman, Wheeler, Wightman, & Carelli, 2008) and NAc neural activity (Delaere, Akaoka, De Vadder, Duchampt, & Mithieux, 2013; Heijkoop, et al., 2024; Stice, Yokum, Blum, & Bohon, 2010), while chronic exposure leads to an attenuated NAc response (Fam, Clemens, Westbrook, Morris, & Kendig, 2022; Giuliani, Mann, Tomiyama, & Berkman, 2014; Heijkoop, et al., 2024) and lower dopamine turnover in the NAc and VTA Davis, et al., 2008; Geiger, et al., 2008; Geiger, et al., 2009; Martel & Fantino, 1996a, 1996b; Nguyen, et al., 2017; Rada, Bocarsly, Barson, Hoebel, & Leibowitz, 2010; South, Westbrook, & Morris, 2012; Totten, Wallace, Pierce, Fordahl, & Erikson, 2022). Chronic junk food exposure also affects D1 and D2 receptors in the NAc (Burke & Small, 2016; Johnson & Kenny, 2010; Robinson, et al., 2015; Steele, et al., 2010; Wang, et al., 2001) and opioid signaling in the VTA (Blendy, et al., 2005; Ong, Wanasuria, Lin, Hiscock, & Muhlhausler, 2013; Robinson, et al., 2015; Smith, Harrold, & Williams, 2002; Vucetic, Kimmel, & Reyes, 2011).

Human studies on direct junk food effects are scarce, but high-fat, high-sugar yogurt consumption diminishes low-fat food preference (Thanarajah, et al., 2023). Furthermore, overweight/obesity is associated with increased sensitivity to immediate rewards, delay-discounting, eating-related self-control, and reward cue tracking (Davis & Fox, 2008; Dietrich, Federbusch, Grellmann, Villringer, & Horstmann, 2014; Horstmann, et al., 2011; Meemken, Kube, Wickner, & Horstmann, 2018; Weller, Cook, Avsar, & Cox, 2008), while inhibitory responses, contingency learning, probabilistic learning, and reward devaluation effects are negatively correlated with overweight and obesity (Coppin, Nolan-Poupart, Jones-Gotman, & Small, 2014; Horstmann, et al., 2015; Janssen, et al., 2017; Kube, et al., 2018; Mathar, Neumann, Villringer, & Horstmann, 2017; Nederkoorn, Smulders, Havermans, Roefs, & Jansen, 2006; van den Akker, Schyns, & Jansen, 2017).

Many rat studies have investigated the direct effects of long-term junk food consumption on behavior but show conflicting behavioral results. While some studies reported increased motivation to work for food rewards (Greenwood, Quartermain, Johnson, Cruce, & Hirsch, 1974; Harb & Almeida, 2014; la Fleur, et al., 2007; Narayanaswami, Thompson, Cassis, Bardo, & Dwoskin, 2013; Robinson, et al., 2015; Vasselli, Cleary, Jen, & Greenwood, 1980), others found no changes or reduced interest in cues for reward (Davis, et al., 2008; Duca, Swartz, & Covasa, 2014; Ducrocq, et al., 2019; Fam, et al., 2022; Shin, Townsend, Patterson, & Berthoud, 2011; Spaulding, et al., 2024; Steele, Pirkle, Davis, & Kirkpatrick, 2019). These inconsistencies are most likely caused by variations in diet types, durations, and models used, along with differences in food rewards.

Interestingly, food rewards that are used in behavioral tests are often different from the palatable diet used as initial manipulation. This suggests that junk food effects extend beyond the original palatable food. It has been shown that long-term junk food exposure could alter the behavioral responses toward other kinds of unnatural drug rewards, like ethanol, amphetamine, cocaine and brain stimulation (Blanco-Gandia, et al., 2017; Brown, Dayas, James, & Smith, 2022; Cook, et al., 2017; Davis, et al., 2008; Gac, et al., 2015; Johnson & Kenny, 2010; Takase, Tsuneoka, Oda, Kuroda, & Funato, 2016). Similarly, human studies link obesity to less efficient monetary reward responses (Balodis, et al., 2013; Kube, et al., 2018; Opel, et al., 2015).

Human fMRI studies have limitations in high-resolution brain investigations or to study diet manipulations consequences. Rat studies, on the other hand, offer a unique opportunity to explore neural regulations in greater detail linked to long-term junk food consumption behaviors. Still, many studies focus only on a small fraction of the reward-related behavior and use operant conditioning designs dependent on learning and motor skills. The impact of junk food consumption on rewarding behavior patterns with free food reward access and interaction remains unknown. Prolonged junk food exposure may affect other rewarding behaviors using the same neural pathways such as sexual behavior, which involves the VTA and NAc (Caggiula, et al., 1979; Guarraci, Megroz, & Clark, 2002; Hasegawa, Takeo, Akitsu, Hoshina, & Sakuma, 1991; Pfaus, Damsma, Wenkstern, & Fibiger, 1995; Rivas & Mir, 1990; Sirinathsinghji, Whittington, & Audsley, 1986). By combining behavioral analysis with real-time VTA neural activity recording, we studied the effects of excessive junk food consumption on eating and sexual behaviors, and the VTA response upon a reward-predictive light cue.

Finally, rodents in junk food exposure studies are often fed high-fat and/or high-sugar pellets, controlling nutrients and energy intake, but not reflecting human dietary behaviors that involve taste, texture, novelty, and variety (McCrickerd & Forde, 2016). Choice and meal variety significantly influence overeating, even in rats (la Fleur, Luijendijk, van der Zwaal, Brans, & Adan, 2014; Rolls, et al., 1981). *Cafeteria diet* (CAF), as used in this study, offers rats *a choice* of the unhealthy, highly palatable human foods (e.g., bacon, muffins, and cookies), enhancing hedonic properties and consumption motivation (Lalanza & Snoeren, 2021; Martire, Holmes, Westbrook, & Morris, 2013; Sampey, et al., 2011; Scheggi, Secci, Marchese, De Montis, & Gambarana, 2013).

Our study investigated whether excessive consumption of junk food, in the form of CAF, alters VTA neural activity patterns in response to food and sexual rewards in rats. We hypothesized that long-term CAF exposure disrupts the VTA response to food reward and in other reward contexts, like pursuing a sexual partner. Moreover, we expected CAF diet to impact brain reward circuitry more severely than the high-energy (HE) pellet diet commonly used in obesity research. Finally, we investigated whether reversal to a healthy diet could restore the behavioral and neural changes caused by long-term CAF exposure.

## 2. Methods

### 2.1 Rats

A total of 36 Wistar female rats (200-250 g upon arrival) and 10 male rats (250 g upon arrival) were purchased from Janvier labs (France). All rats were pair-housed in Macrolon IV ® cages on a reversed 12 h light/dark cycle (lights on between 23:00 and 11:00) in a room with controlled temperature (21 ± 1 °C) and humidity (55 ± 10 %), with ad libitum access to standard rodent food pellets (low phytoestrogen maintenance diet, #V1554, Ssniff, Germany) and tap water throughout the experiment (unless otherwise indicated). All animal care and experimental procedures employed in this study were conducted in agreement with the European Union council directive 2010/63/EU. All animal use was reviewed and approved by the Norwegian Food Safety Authority, # 19682, 8^th^ of July 2020.

### 2.2 Experimental design

Following the acquisition of sexual experience (see supplementary methods for more details), female rats underwent stereotaxic brain surgery in two batches of 18 rats, with two week intervals between batches. After recovery from surgery, the females were randomly assigned to one of three experimental groups: control (CTR), high energy (HE), or cafeteria (CAF) diet for six weeks.

After six weeks of diet exposure, the females were tested in the behavioral reward box (Fig 1A/B), during which they were tethered to the fiber photometry system (RZ10x from Tucker-Davis Technologies, USA). The testing sequence consisted of five daily tests with a food reward over five consecutive days, followed by three tests with a sexual reward on the same day of their proestrus (day of sexual receptivity) to prevent data loss due to pseudopregnancy; a 2-hour break was maintained between tests. The food reward consisted of small portions of the preferred products (as in most consumed) during the CAF diet exposure (Table 1), and a male rat served as the sexual reward.

**Figure 1.**
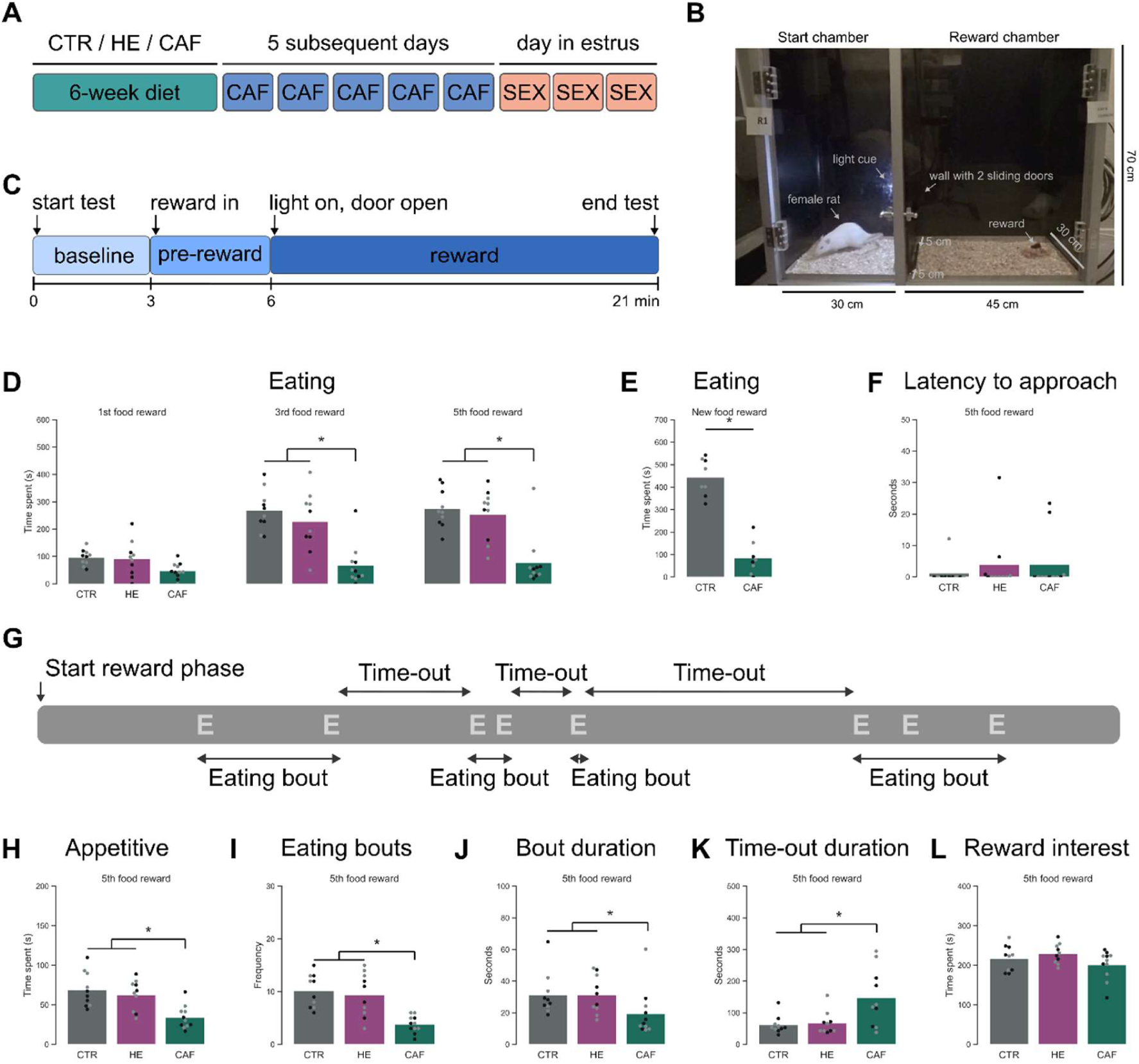
The effects of Cafeteria, High energy, and control diet on behavioral responses during a food reward test. **A)** Schematic representation of the experimental design. **B)** Picture of our behavioral reward test box. **C)** Schematic representation of the behavioral reward test. **D)** Time spent eating **E)** Time spent eating a novel CAF reward. CTR: n=8, CAF: n=8. **F)** Latency to interact with the CAF reward. **G)** Schematic example of eating bouts and time-outs. E = eating **H)** Number of appetitive behaviors. **I)** Number of eating bouts. **J)** Mean duration of eating bouts **K)** Mean duration of time-outs. **L)** Time spent on behaviors that indicate interest in the reward. **All** figures show rats on control (CTR, grey bar), high energy pellet (HE, purple bar), and cafeteria (CAF, green bar) diet during the reward phase of the 1^st^, 3^rd^, and/or 5^th^ food reward test, unless stated differently. Dot colors representing their body weight category (grey: lighter than average, black: heavier than average). CTR n=10, HE n=10, CAF n=11. * p<0.05 compared to by line connected group.

**Table 1:**
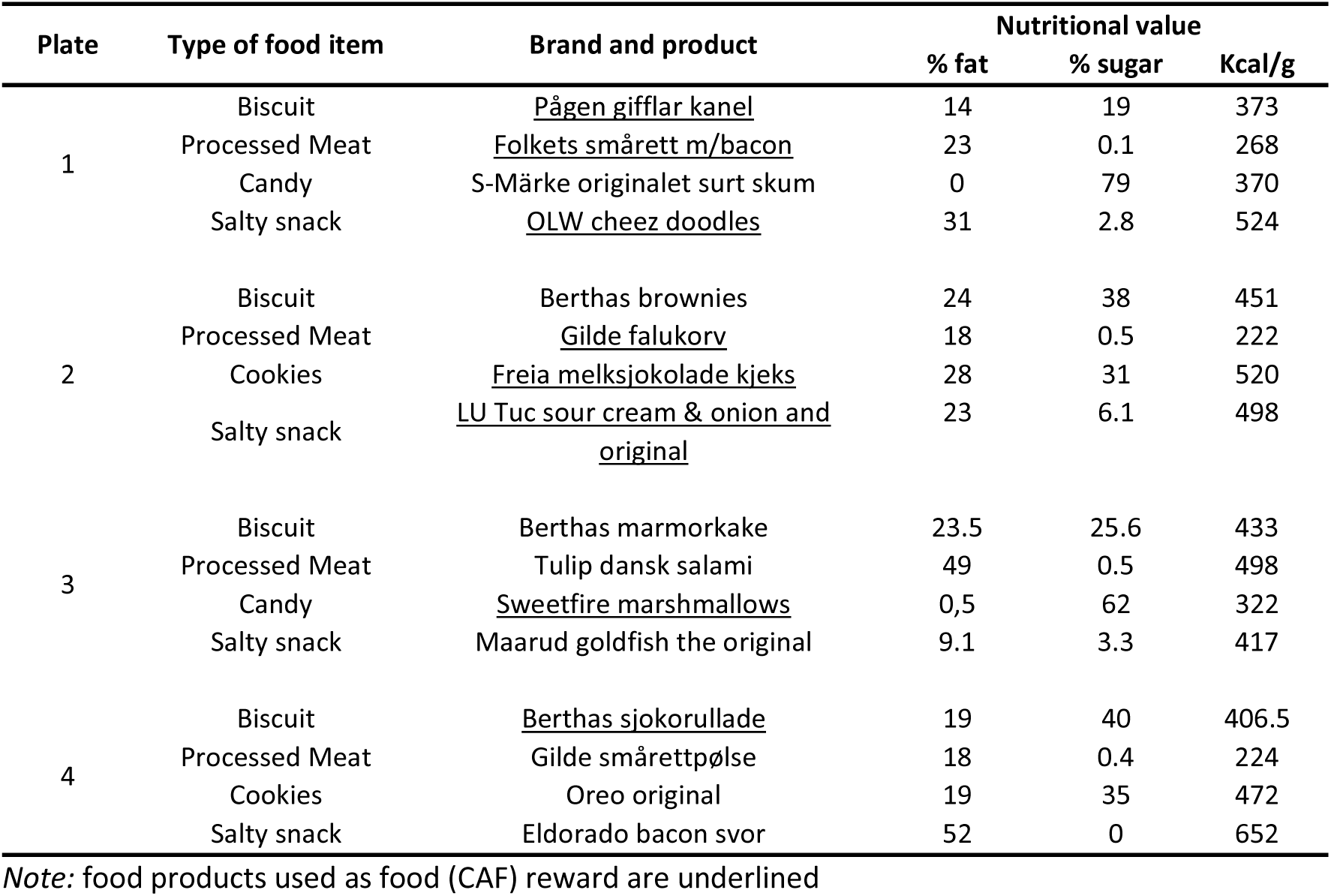
Food items for each of the 4-plates of the Cafeteria diet (CAF)

To mitigate the potential effects of pseudopregnancies, the female rats remained on their experimental diet for an additional two weeks following the behavioral reward tests. Subsequently, all female rats were switched to a standard chow diet for another two weeks (reversal diet). This final testing phase included three consecutive daily tests with the food (CAF) reward and one test with the sexual reward on a day in estrus.

After completion of the experiment, all rats were perfused and their brains were collected to assess viral infusion location and expression and verification of fiber location. See supplementary methods for more details on brain processing, immunostaining and imaging.

### 2.3 Surgeries

#### 2.3.1 Brain surgery

All female rats were anesthetized with isoflurane, placed in a stereotaxic apparatus (Stoelting Europe, Ireland), and received 0.05 mg/kg buprenorphine and 2 mg/kg meloxicam subcutaneously. The scalp was shaved, disinfected with povidone-iodine and locally anaesthetized with bupivacaine (0.15 ml, s.c.). Core body temperature, oxygen saturation and heart rate were monitored throughout the surgery. A hole was drilled above the VTA of the right hemisphere (coordinates: AP -5.0 mm, ML +0.9 mm, DV -8.5 mm relative to bregma (Paxinos & Watson, 1998), selected based on (Beloate, Omrani, Adan, Webb, & Coolen, 2016)) and a 2 µl Hamilton Neuros syringe with a 30G blunt needle mounted in a Hamilton injector (Stoelting Europe, Ireland) on the stereotaxic apparatus was slowly inserted. N=11 of each group was infused with 750 nl, 1×10¹³ vg/mL pAAV5.syn.GCaMP6s.WPRE.SVA40 (a gift from Douglas Kim & GENIE Project (Addgene, Massachusetts, USA, plasmid # 100843 ; http://n2t.net/addgene:100843; RRID:Addgene_100843-AAV5) to induce the expression of GCaMP6s, and N=1 of each group with 750 nl, 1×10¹³ vg/mL pAAV5.hSyn.eGFP.WPRE.bGH (a gift from James M. Wilson (Addgene Massachusetts, USA, plasmid # 105539 ; http://n2t.net/addgene:105539 ; RRID:Addgene_105539-AAV5)) to induce the expression of eGFP as control, both at a rate of 125 nl/min. Following infusion, the needle was left in place for 10 minutes before withdrawal. 4 anchor screws (1 x 3 mm, Agnthos, Sweden) were placed in the skull, after which the fiber-optic cannula (diameter 400 µm, NA 0.66, Doric Lenses, Quebec, Canada) was inserted in the right hemisphere at AP -5.0 mm, ML +0.9 mm, DV -8.4 mm relative to bregma. It was fixed to the skull with super bond (C&B Super bond, Technomedics, Norway) followed by regular dental acrylic (Meliodent, Kulzer GmbH, Germany). Post-operative analgesic treatment consisted of 0.05 mg/kg buprenorphine and 2 mg/kg meloxicam after 24 and 48 hours. Immediately after surgery the rats were pair-housed and allowed a minimum of one week before initiation of the diet manipulation.

#### 2.3.2 Vasectomy

To prevent pregnancies in experimental female rats while maintaining their reproductive integrity, the stimulus male rats were subjected to vasectomy by ligation of a small portion of the vas deferens of both testes, as previously described (Phang, 1993). On each side, a small incision was made in the scrotal skin covering the testes and the tissue layer covering the testicle was opened. The testicle, epididymis and vas deferens were exposed and the vas deferens was thermally cauterized. The organs were returned inside the incision and the wound was closed with stitches. Immediately after surgery the male rats were pair-housed and were allowed to recover for two weeks before obtaining sexual experience.

### 2.4 Diet manipulation

Experimental rats were randomly assigned to one of three experimental diets for a period of six weeks:

- Standard Chow (CTR): standard laboratory chow.
- High energy (HE): laboratory chow with high content of fat and sugar that simulates the nutritional values of junk food, but in the shape of regular pellets (Western Diet #D12079B, Research Diets Inc., USA).
- Cafeteria diet (CAF): composed of standard chow and 16 different food items obtained in a supermarket, including biscuits, cookies, processed meat and salty snacks (Table 1 and S1), based on a proposed protocol for cafeteria diet (Lalanza & Snoeren, 2021). The food items were divided into 4-daily-rotated plates balanced by type of food, texture, taste, and nutritional composition.

All three experimental diets were provided *ad libitum* to the rats, along with access to tap water *ad libitum*.

### 2.5 Behavioral testing

All rats underwent a 10-minute habituation to the behavioral reward test box (see supplementary methods for more details on the box) on the day preceding commencement of testing. On the day of behavioral testing, food was removed from the cage at 09:00, while access to water remained available. Behavioral testing was conducted between 12:00 and 19:00, after which fresh food was returned to the cage. The diet groups were randomly distributed across the testing period and no order effects were detected.

First, a subject rat was connected to the fiber photometry system (see supplementary methods for details on fiber photometry recording and processing) using a fiber-optic cable and placed in a small cage for approximately 2 minutes to allow the GCaMP and isosbestic signals to stabilize. The reward test started when the female subject was placed in the left chamber, with the doors in the dividing wall closed (Baseline phase, Fig 1C). After 3 minutes, a reward (either food or a male rat) was placed in the right chamber, while the doors remained closed for another 3 minutes (Pre-reward phase).

Subsequently, the light turned on for 10 seconds, after which the doors opened. The light turned off again, and the rat was allowed to interact freely with the reward for a total of 15 minutes (Reward phase). The doors opened to a width of 5 cm, allowing the females to cross easily between compartments, while males spent more time crossing due to their larger body sizes. This configuration enabled the females to pace their sexual activity slightly more than with completely open doors. After the test, the female was returned to the home cage, and the reward box was cleaned with ethanol. Fresh bedding was provided for the next rat.

All behavioral reward tests were recorded on video with a digital video camera (Sony HDR-CX240E handycam) for behavioral assessment.

### 2.6 Behavioral assessment

From the videos, which were recorded at a frequency of 25 fps, the frequency and duration of the multiple behaviors (Table 2) were annotated, and the start of the behaviors was time-locked on the exact video frames using The Observer XT12.5 and Observer XT16 (Noldus, The Netherlands).

**Table 2:** Description of behaviors scored in the different behavioral reward tests.

| Behavior | Description |
| --- | --- |
| Approaching reward | Intentional and direct movement (approach) towards the reward. |
| Sniffing reward | Sniffing the reward actively. |
| Carrying food | Touching, holding or moving the reward. |
| Eating | Eating the reward. |
| Darts | Fast runaway behavior ending with a sudden stop allowing the male to mount (part of paracopulatory behavior). |
| Hops | Short jumping behavior with all legs off the ground (part of paracopulatory behavior). |
| Anogenital sniffing | Sniffing the anogenital region of the stimulus male (sexual reward) |
| Lordosis | Receptive behavior in which the female rat arches her back and deflects her tail to one side allowing the male access to her vagina. |
| Lordosis score | The intensity of the lordosis responses scored from 0 (no lordosis upon a sexual stimulation) to 3 (exaggerated lordosis) (Hardy & DeBold, 1971). Lordosis score is calculated by number of lordosis 1 * 1 + number of lordosis 2 * 2 + number of lordosis 3 * 3) / number of received copulations. |
| Lordosis quotient | The number of lordosis responses (including those without received copulations) divided by the number of received copulations. |
| Received copulations | Male's sexual behavior including *mount*: pelvic thrusting while being mounted on the female; *intromission*: mounting the female with pelvic thrusting and penile insertion into the vagina; characterized by a more vigorous dismount; *ejaculation*: characterized by a longer intromission with extra muscles contraction during climax, and the female moving away from the male. |
| Eating bout | One or multiple eating episodes that were not interrupted by any behavior that was not related to the food reward. As soon as a behavior unrelated to the food reward was displayed, the last eating episode was considered the end of the sexual bout. |
| Sexual bout | One or multiple episodes containing paracopulatory and lordosis behavior that were not interrupted by any behavior that was not related to the sexual reward. As soon as a behavior unrelated to the sexual reward was displayed, the last paracopulatory or lordosis episode was considered the end of the sexual bout. |
| Time-out | Time elapsed from the end of an eating or sexual bout to the start of the next eating or sexual bout. |
| Exploring environment | Active exploration of the chambers such as rearing or sniffing the bedding and walls. |
| Exploring door | Active interaction with the inner door, such as rearing or sniffing the door. |
| Self-grooming | Cleaning behavior of the body and face. |
| Head away from reward | Turning the head (and thus attention) away from the reward |
| Head towards reward | Turning the head (and thus attention) towards the reward |

Then, some additional parameters were calculated, such as the lordosis score ((number of lordosis with score 1 times 1 + number of lordosis with score 2 times 2 + number of lordosis with score 3 times 3) / number of received copulations) (Hardy & Debold, 1971), lordosis quotient (number of lordosis responses (1-3) / number of received copulations * 100%) , and the mean contact-return latency after a received mount, intromission or ejaculation (latency to 1^st^ paracopulatory behavior after receiving a copulation) were calculated per rat.

Moreover, a more detailed behavioral assessment paradigm was used to explore micropatterns in reward-related behaviors. For this analysis, behavior was divided into eating bouts (Fig 1G, food reward tests) and sexual bouts (Fig 3F, sexual reward test). An eating bout was defined as all eating episodes uninterrupted by any behavior that was not focused on the food reward. As soon as a behavior not directed at the food reward was displayed, the last eating episode was considered the end of the eating bout. The interval until the next eating episode was termed a time-out. A similar approach was applied to define a sexual bout, where both lordosis and paracopulatory behaviors could initiate and/or terminate a sexual bout. The number of bouts, the mean duration of bouts, and the mean time-out duration were quantified. (Note that eating bouts are different from previously used meal-analyses (Ali & Kravitz, 2018; Zorrilla, et al., 2005).)

To mitigate the effects of potential pseudopregnancies, all three sexual reward tests were conducted on a single day and with the same male for each female rat. Regrettably, not all females reached the estrus phase and were consequently excluded from the analysis. Notably, only six out of ten females on the HE diets reached estrus, and two of these females did not engage in copulation with the male. Furthermore, although ten of our CAF-exposed females reached the estrus phase, three of them did not exhibit any paracopulatory behaviors during the first sexual reward test. This observation suggests a potential delay in the onset of the receptive phase compared to CTR females.

To address these challenges in conducting a reliable behavioral reward test with a sexual reward, we analyzed the neural and behavioral data from the most successful sexual reward test. The *best sexual reward test* was defined as the test (either the first, second, or third) in which the female displayed the highest number of paracopulatory behaviors. In practice this meant that two 1^st^, four 2^nd^ and four 3^rd^ tests were included in the analysis for the CTR rats; one 1^st^, two 2^nd^ and three 3^rd^ tests for the HE rats; and two 1^st^, four 2^nd^, and three 3^rd^ tests for the CAF rats. The sex reward test after reversal was conducted later on the days (2.5 hours after the lights went off) to avoid the delay in the onset of the receptive phase.

### 2.7 Statistical analysis

All rats were included in the behavioral analysis, except for three rats that were excluded due to the loss of their fiber caps at an early stage of the experiment.

For the fiber photometry analysis, seven rats were excluded from analysis due to lack of GCaMP6s expression or incorrect fiber placement, resulting in the inclusion of 6 CTR-, 7 HE- and 9-CAF included rats. In addition, a small number of sessions (1 CTR best sexual reward tests and 1 CAF sexual reward test after the reversal diet) were excluded from the analysis due to insufficient quality of the fiber photometry recording.

Both the behavior and fiber photometry data were analyzed with customized Python 3.9 scripts (available upon request). Statistical analysis of the behavioral data and area under the curve were conducted using SPSS software (version 29, IBM, Armonk, USA) with a level for statistically significant difference set at p < 0.05. For the behavioral analysis in which CTR, CAF, and HE were compared, a linear mixed model was used to investigate the diet, test and diet*test interaction effects. In the case of a significant effect, Bonferroni-corrected post hoc t-tests were conducted. When CTR rats were compared to CAF rats only, an independent sample t-test was used. The linear mixed model analyzing the AUC data comprised the factors Diet (CTR, HE, and CAF or CTR and CAF) and Epoch (pre versus post). The pre vs post effects on AUC for the light and door cue were analyzed separately. In case the test was divided into 3 parts, the factor Parts (part 1, part 2, and part 3) was also included in the analysis. Again, in the case of a significant effect, Bonferroni-corrected post hoc t-tests were conducted.

## 3. Results

To study the impact of long-term CAF diet exposure on the VTA during behavioral responses to food or sexual rewards, we assigned adult female Wistar rats to one of three dietary groups: a control diet (CTR, standard chow), a high energy diet (HE, RD Western Diet pellets), or a cafeteria diet (CAF, see Table 1 for the details of the products, Table 3 for the details of the body weights, and tables S1-S3 for more detail about the food intake and nutritional values per cages). After 6-weeks of diet exposure, rats underwent a sequence of behavior tests with a food reward (consisting of a selection of the preferred CAF diet items), followed by a sexual reward (a sexually trained male rat, Fig 1A/B). Our experimental design facilitated the assessment of distinct phases of reward-related behavior, including a baseline phase (without any reward), a pre-reward phase (introducing the reward without access), and a reward phase allowing free interaction with the reward (Fig 1C).

**Table 3:** Body weights and body weight gains per rat.

| Cage | Rat | Diet | Body weight (g) week 1 | Body weight (g) week 7 | Body weight gain (g) |
| --- | --- | --- | --- | --- | --- |
| 1 | F301 | CAF | 245 | 318 | 74 |
|  | F302 | CAF | 259 | 320 | 61 |
| 2 | F303 | CTR | 243 | 277 | 34 |
|  | F304 | CTR | 305 | 304 | -1 |
| 3 | F305 | HE | 290 | 329 | 39 |
|  | F306 | HE | 252 | 292 | 40 |
| 4 | F307 | HE | 256 | 309 | 53 |
|  | F308 | HE | 258 | 291 | 33 |
| 5 | F309 | CAF | 253 | 303 | 50 |
|  | F310 | CAF | 216 | 260 | 44 |
| 6 | F311 | CTR | 254 | 287 | 34 |
|  | F312 | CTR | 265 | 289 | 25 |
| 7 | F313 | CAF | 318 | 309 | -9 |
|  | F314 | CAF | 255 | 307 | 52 |
| 8 | F315 | CTR | 268 | 300 | 32 |
|  | F316 | CTR | 268 | 282 | 14 |
| 9 | F317 | HE | 272 | 335 | 63 |
|  | F318 | HE | 264 | 314 | 50 |
| 10 | F319 | CAF | 235 | 291 | 56 |
|  | F320 | CAF | 285 | 355 | 70 |
| 11 | F321 | HE | 292 | 340 | 48 |
|  | F325 | HE | 271 | 314 | 43 |
| 12 | F323 | CTR | 276 | 298 | 22 |
|  | F324 | CTR | 273 | 289 | 16 |
| 14 | F327 | CTR | 255 | 271 | 16 |
|  | F328 | CTR | 270 | 256 | -14 |
| 15 | F329 | CAF | 255 | 309 | 54 |
|  | F330 | CAF | 264 | 319 | 55 |
| 16 | F331 | CAF | 257 | 302 | 45 |
|  | F332 | CAF | 253 | 326 | 73 |
| 17 | F333 | HE | 278 | 315 | 37 |
|  | F334 | HE | 267 | 308 | 41 |
| 18 | F335 | CTR | 276 | 302 | 26 |
|  | F336 | CTR | 254 | 268 | 14 |
|  |  | CTR | 268.5 ± 5.32 | 283.6 ± 4.98 | 15.1 ± 4.38 |
|  |  | CAF | 252.5 ± 5.18 | 310.0 ± 7.08* | 57.5 ± 3.19* |
|  |  | HE | 270.0 ± 4.28 | 314.7 ± 5.18* | 44.7 ± 2.82*^ |

Note 1: \* p<0.05 compared to CTR, ^ p<0.05 compared to CAF.

### 3.1 Long-term CAF diet exposure, in contrast to HE, reduces interest in a food reward

First, we investigated whether long-term CAF diet exposure altered behavioral responses towards a food reward. We found that CAF-exposed rats exhibited a reduced frequency of eating (Fig S1A, Diet: F_(2,28)_=25.121, p<0.001, Test: F_(2,56)_=30.493, p<0.001, Diet*Test: F_(4,56)_=5.734, p<0.001, post-hoc: 3^rd^ test: CAF vs HE: p<0.001, CAF vs CTR: p<0.001, 5^th^ test: CAF vs HE: p<0.001, CAF vs CTR: p<0.001, CTR: 1^st^ vs 3^rd^ test: p<0.001, 1^st^ vs 5^th^ test: p<0.001, HE: 1^st^ vs 3^rd^ test: p<0.001, 1^st^ vs 5^th^ test: p<0.001 (note: only the significant post-hoc results are reported in the result section) and spent notably less time consuming the reward compared to both CTR and HE rats (Fig 1D, Diet: F_(2,28)_=29.791, p<0.001, Test: F_(2,56)_=28.911, p<0.001, Diet*Test: F_(4,56)_=4.754, p=0.002) during the reward phase. This disparity became apparent in subsequent tests as HE and CTR rats increased their time spent eating, while CAF rats did not. (3^rd^ test: CAF vs HE: p<0.001, CAF vs CTR: p<0.001, 5^th^ test: CAF vs HE: p<0.001, CAF vs CTR: p<0.001, CTR: 1^st^ vs 3^rd^ test: p<0.001, 1^st^ vs 5^th^ test: p<0.001, HE: 1^st^ vs 3^rd^ test: p<0.001, 1^st^ vs 5^th^ test: p<0.001).

To investigate the potential influence of novelty on reward consumption, we repeated the study with CTR and CAF rats, introducing a novel CAF reward (with human junk food items they had never eaten before). CAF rats spent less time eating the novel reward compared to CTR rats (Fig 1E, *t*-test: *t*=-8.78, p<0.001), suggesting that novelty did not account for the reduced consumption in CAF rats.

Further analysis revealed that the latency to approach food rewards was similar across all diet groups in all tests (Fig 1F, 5th test, Diet: F_(2,28)_=0.987, p=0.385, Test: F_(2,56)_=0.733, p=0.485, Diet*Test: F_(4,56)_=0.786, p=0.539). However, CAF rats spent less time sniffing and approaching the food post-initial approach (appetitive behavior, Fig 1H, 5th test, Diet: F_(2,28)_=16.296, p<0.001, Test: F_(2,56)_=6.666, p=0.003, Diet*Test: F_(4,56)_=3.010, p=0.026, post-hoc: 3^rd^ test: CAF vs HE: p=0.002, CAF vs CTR: p<0.001, 5^th^ test: CAF vs HE: p=0.003, CAF vs CTR: p<0.001, CTR: 1^st^ vs 3^rd^ test: p<0.001, 1^st^ vs 5^th^ test: p=0.074, HE: 1^st^ vs 3^rd^ test: p=0.033). CAF rats also showed a decrease in number of appetitive behaviors across tests, while CTR rats increased these behaviors (Fig S1B, Diet: F_(2,28)_=12.382, p<0.001, Test: F_(2,56)_=4.381, p=0.017, Diet*Test: F_(4,56)_=11.795, p<0.001, post-hoc: 3^rd^ test: CAF vs HE: p<0.001, CAF vs CTR: p<0.001, 5^th^ test: CAF vs HE: p<0.001, CAF vs CTR: p<0.001, HE vs CTR: p=0.051, CAF: 1^st^ vs 3^rd^ test: p=0.024, 1^st^ vs 5^th^ test: p<0.001, CTR: 1^st^ vs 3^rd^ test: p=0.001, 1^st^ vs 5^th^ test: p<0.001, HE: 1^st^ vs 3^rd^ test: p<0.001).

In contrast to more traditional conditioning tests, providing the rats with continuous free access to the reward allowed for a detailed behavioral assessment. We defined eating bouts as a sequence of eating episodes uninterrupted by non-reward-related behaviors (Fig. 1G, similar approach as previous studies on sexual behavior (Huijgens, Guarraci, Olivier, & Snoeren, 2021; Huijgens, Heijkoop, & Snoeren, 2021; Huijgens, Heijkoop, Vanderschuren, Lesscher, & Snoeren, 2024; Oyem, Heijkoop, & Snoeren, 2025)). Our findings revealed that CAF rats exhibited fewer and shorter eating bouts (frequency: Fig 1I and Fig S1C, Diet: F_(2,28)_=17.893, p<0.001, Test: F_(2,56)_=26.634, p<0.001, Diet*Test: F_(4,56)_=5.287, p<0.001, post-hoc: 3^rd^ test: CAF vs HE: p<0.001, CAF vs CTR: p<0.001, 5^th^ test: CAF vs HE: p<0.001, CAF vs CTR: p<0.001, CTR: 1^st^ vs 3^rd^ test: p<0.001, 1^st^ vs 5^th^ test: p<0.001, HE: 1^st^ vs 3^rd^ test: p<0.001; mean duration: Fig 1J and Fig S1D, Diet: F_(2,27.981)_=7.622, p=0.002, Test: F_(2,54.709)_=5.478, p=0.007, Diet*Test: F_(4, 54.709)_=1.321, p=0.274, post-hoc: 3^rd^ test: CAF vs CTR: p=0.023, 5^th^ test: CAF vs HE: p=0.029, CAF vs CTR: p=0.029, CTR: 1^st^ vs 3^rd^ test: p=0.016, 1^st^ vs 5^th^ test: p=0.005), with longer time-outs between the bouts (Fig 1K and Fig S1E, Diet: F_(2,27.327)_=8.978, p=0.001, Test: F_(2,53.157)_=8.354, p<0.001, Diet*Test: F_(4, 53.141)_=0.492, p=0.741, post-hoc: 5^th^ test: CAF vs HE: p=0.005, CAF vs CTR: p=0.003, HE: 1^st^ vs 3^rd^ test: p=0.026) compared to CTR and HE rats.

No differences were found in baseline activity, with all rats spending similar amount of time exploring the chamber (Fig S1F, Diet: F_(2,28)_=1.705, p=0.200, Test: F_(2,56)_=6.395, p=0.003, Diet*Test: F_(4, 56)_=0.879, p=0.483, post-hoc: CAF: 1^st^ vs 5^th^ test: p=0.031, CTR: 1^st^ vs 5^th^ test: p=0.050) and engaging in grooming activities (Fig S1G, Diet: F_(2,28)_=0.264, p=0.770, Test: F_(2,56)_=0.268, p=0.766, Diet*Test: F_(4, 56)_=0.801, p=0.529). During the pre-reward phase, CAF rats did not differ in anticipatory behaviors (such as exploring the closed door) compared to HE and CTR rats (Fig 1L, Diet: F_(2,28)_=3.195, p=0.056, Test: F_(2,56)_=10.240, p<0.001, Diet*Test: F_(4, 56)_=1.186, p=0.327, post-hoc: CTR: 1^st^ vs 5^th^ test: p=0.008, HE: 3^rd^ vs 5^th^ test: p=0.028). However, as expected, all rats spent approximately twice as much time engaging in these behaviors during the pre-reward phase compared to the baseline phase when no reward was present behind the doors (Fig 1L compared to Fig S1H, Diet: F_(2,28)_=1.622, p=0.216, Test: F_(2,56)_=2.737, p=0.073, Diet*Test: F_(4, 56)_=0.357, p=0.838).

Overall, these findings suggest that long-term exposure to a CAF diet alters both appetitive and consummatory behavioral responses to a food reward, while anticipatory behaviors and reward-unrelated behaviors remain unaffected. Notably, long-term exposure to a HE diet did not produce the same effects, resulting in comparable levels of behavioral interest in the food reward as observed in CTR rats.

### 3.2 Long-term CAF diet exposure reduces VTA activity in response to food rewards

To investigate the impact of long-term CAF diet exposure on VTA neural activity during reward responses, all rats underwent stereotaxic surgery to induce GCaMP6s expression in VTA neurons (Fig 2A). Following a one-week recovery period, rats were placed on their respective diets for six weeks, after which (the above-mentioned) behavioral testing commenced and VTA activity was recorded.

**Figure 2.**
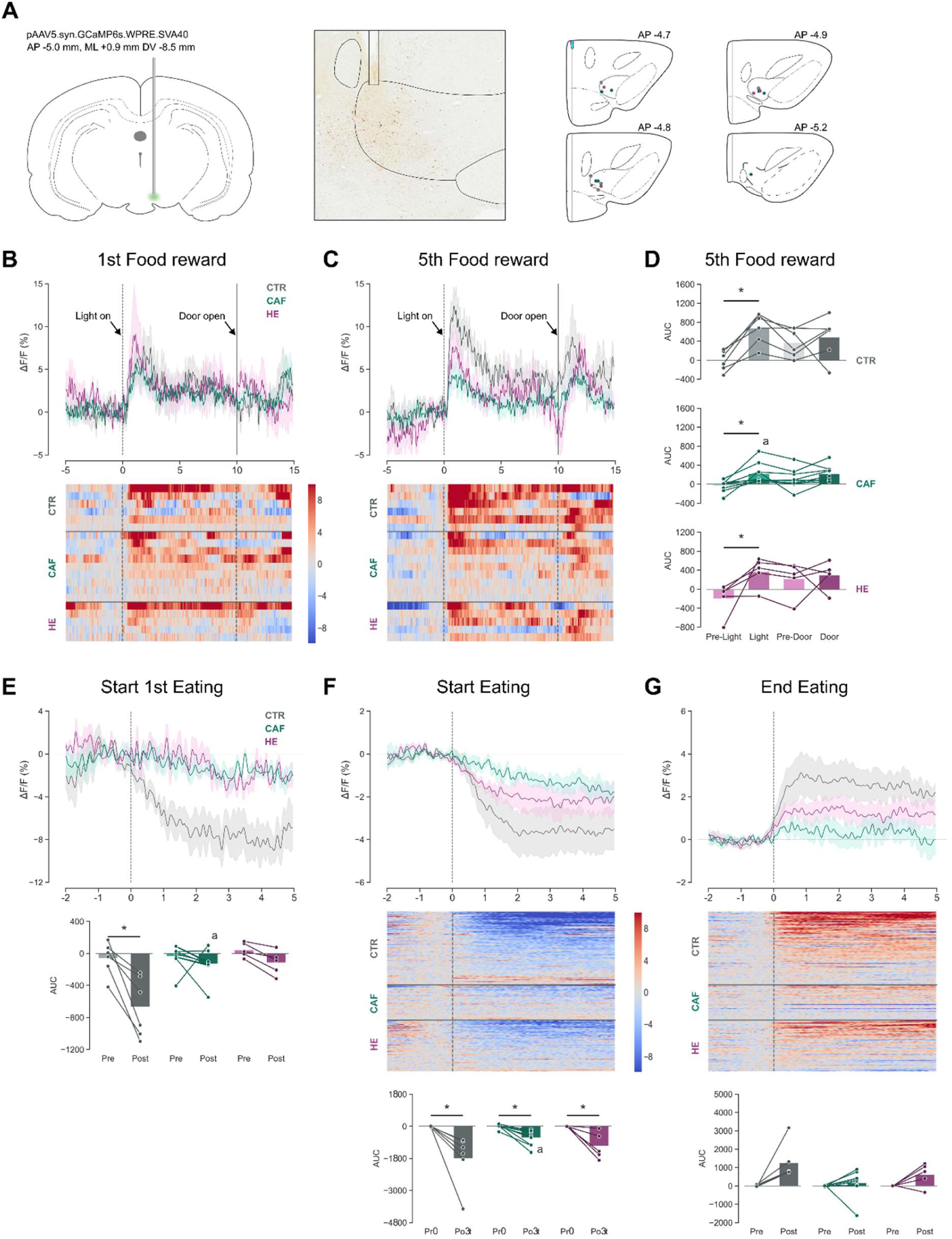
The effects of Cafeteria, High fat high sugar, and control diet on the ventral tegmental area (VTA) during a food reward test. **A)** Schematic representation of surgical details and histologically determined fiber locations and AAV vector expression in the VTA. **B)** Upper panel: ΔF/F aligned to the light cue at t=0 and door opening at t=10s. Lower panel: heat maps of ΔF/F aligned to the light cue and door opening. **C)** Upper panel: ΔF/F aligned to the light cue and door opening. Lower panel: heat maps of ΔF/F aligned to the light cue and door opening. **D)** AUC before (pre-Light) and after (Light) the light cue, and before (Pre-door) and after (Door) the door opening. **E)** Upper panel: ΔF/F aligned to the start of eating episodes. Lower panel: AUC before (pre) and after (post) the start of the very first eating episodes in the 5th reward test with a food reward. **F)** Upper panel: ΔF/F aligned to the start of eating episodes. Middle panel: heat maps of ΔF/F aligned to the start of eating episodes. Lower panel: AUC pre and post the start of eating episodes. **G)** Upper panel: ΔF/F aligned to the end of eating episodes. Middle panel: heat maps of ΔF/F aligned to the end of eating episodes. Lower panel: AUC pre and post the end of eating episodes. **All** Figures show rats on control (CTR, grey color bars and lines) and cafeteria (CAF, green color bars and lines) diets during the reward phase of all behavioral reward tests with a food reward, unless stated otherwise. The dots in the behavioral data have colors representing their body weight category (grey: lighter than average, black: heavier than average). The lines in the ΔF/F figures represent the mean and shaded areas the SEM. CTR n=6, CAF n=9, HE n=5. * p<0.05 compared to by line connected group. ^a^ p<0.05 compared to same bar of CTR.

**Figure 3.**
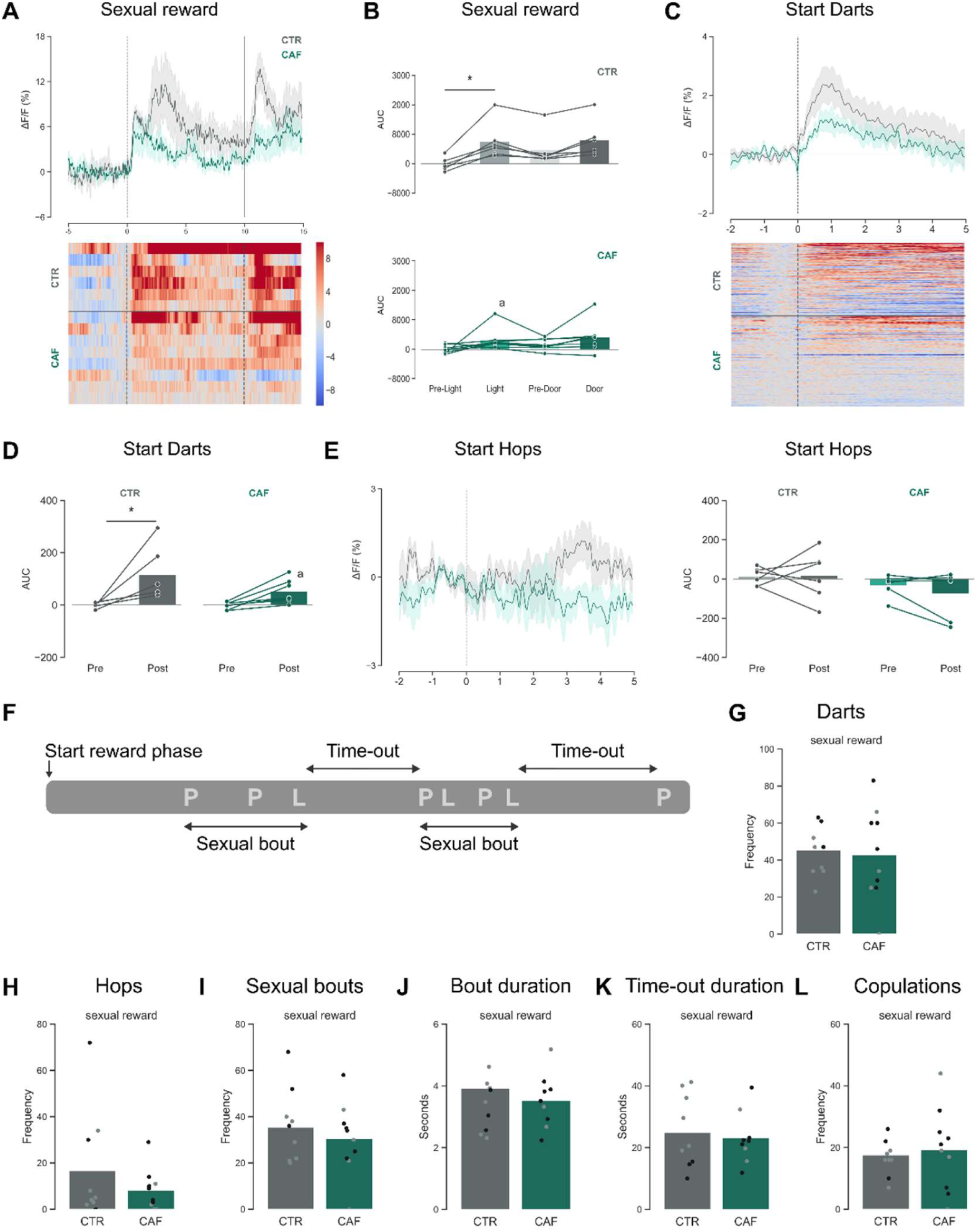
The effects of Cafeteria diet on the ventral tegmental area and behavioral responses during a sexual reward test. **A)** Upper panel: ΔF/F aligned to the light cue at t=0 and door opening at t t=10. Lower panel: heat maps of ΔF/F aligned to the light cue and door opening. **B)** AUC before (pre-Light) and after (Light) the light cue, and before (Pre-door) and after (Door) the door opening. **C)** Upper panel: ΔF/F aligned to the start of darting behaviors. Lower panel: Heat maps of ΔF/F aligned to the start of darting behaviors. **D)** AUC before (pre) and after (post) the start of darting behaviors. **E)** Left panel: ΔF/F aligned to the start of hopping behaviors. Right panel: AUC pre and post the start of hopping behaviors. **F)** Schematic example of sexual bouts and time-outs. P= paracopulatory behaviors, L= lordosis responses. **G)** Total number of darts. **H)** Total number of hops**. I)** Total number of sexual bouts. **J)** Mean duration of a sexual bout. **K)** Mean duration of time-outs. **L)** Total number of copulations received. **All** Figures show rats on control (CTR, grey color bars and lines) and cafeteria (CAF, green color bars and lines) diets during the reward phase of the best sexual reward test. The dots in the behavioral data have colors representing their body weight category (grey: lighter than average, black: heavier than average). The lines in the ΔF/F figures represent the mean and shaded areas the SEM. CTR n=6, CAF n=8 for fiber photometry data and CTR n= 10, CAF n=9 for behavioral data. * p<0.05 compared to by line connected group. ^a^ p<0.05 compared to same bar of CTR.

First, we explored the effects of long-term CAF diet exposure on the VTA in response to cues paired with reward access. During the initial food reward test, an increase in VTA neuronal activity was observed in all rats in response to a light cue signaling the imminent opening of the doors after a 10-second delay (Fig 2B). However, the actual opening of the doors did not elicit a change in VTA activity. Heatmap analysis revealed that the increase in light cue-associated VTA activity was only observed in a selection of rats and varied across dietary groups (Fig 2B). As rats became familiar with the behavioral reward test (on the 5^th^ test day with a food reward), a surge in VTA activity in response to the light cure was observed in CTR rats (Fig 2C), whereas only a subset of HE and CAF rats displayed a similar response. Further analysis of the area under the curve (AUC) revealed that CAF rats, but not HE rats, displayed an attenuated response compared to CTR rats (Fig 2D, Light: Diet: F_(2,34)_=4.591, p=0.017, Epoch: F_(1,34)_=38.576, p<0.001, Diet*Epoch: F_(2, 34)_=2.976, p=0.064, post-hoc: Pre-light vs Light: CTR: p<0.001, CAF: p=0.026, HE: p=0.002, Light: CAF vs CTR: p=0.003, Door: Diet: F_(2,34)_=3.364, p=0.046, Epoch: F_(1,34)_=1.071, p=0.308, Diet*Epoch: F_(2, 34)_=0.023, p=0.977)

Analysis of VTA responses during specific behaviors revealed that CTR rats show a significant reduction in VTA activity (relative to baseline) upon the initial consumption of the reward (Fig 2E, AUC: Diet: F_(2,32)_=3.399, p=0.046, Epoch: F_(1,32)_=7.074, p=0.012, Diet*Epoch: F_(2, 32)_=2.762, p=0.078, post-hoc: Pre vs Post: CTR: p=0.002, Post: CAF vs CTR: p=0.005). In contrast, long-term exposure to HE and CAF was not associated with any change in activity upon this 1^st^ eating episode. Furthermore, CAF rats displayed a pronounced attenuation of ΔF/F event-related transients in the VTA compared to CTR rats when eating the reward, while HE rats had an intermediate response (Fig 2F, AUC: Diet: F_(2,32)_=2.956, p=0.066, Epoch: F_(1,32)_= 31.102, p<0.001, Diet*Epoch: F_(2, 32)_=2.778, p=0.076, post-hoc: Pre vs Post: CTR: p<0.001, HE: p=0.010, CAF: p=0.045, Post: CAF vs CTR: p=0.005).

The decrease in VTA activity was sustained throughout eating episodes. When aligning the fiber photometry signal to the end of each episode of eating, a return to baseline was evident in CTR rats (Fig 2G, shown as an increase due to ΔF/F normalization to baseline at t = 0, AUC: Diet: F_(2,32)_=0.723, p=0.493, Epoch: F_(1,32)_= 0.685, p=0.414, Diet*Epoch: F_(2, 32)_=0.535, p=0.591). Alterations in VTA neural activity upon eating persisted throughout the behavioral reward test. CTR rats consistently displayed activity decreases in the VTA upon eating during the initial, middle, and final five-minute intervals of the test (Fig S2A), whereas CAF rats demonstrated an attenuated response in all intervals. This effect was also reflected in the AUC analysis, with CAF rats showing a significant attenuation compared to CTR rats in all test intervals (Fig S2B, AUC: Diet: F_(2,100)_=7.104, p=0.001, Epoch: F_(1,100)_= 85.933, p<0.001, Part: F_(2,100)_= 0.644, p=0.528, Diet*Epoch*Part: F_(12, 100)_=1.429, p=0.165, post-hoc: pre vs post: CTR: part 1: p<0.001, part 2: p<0.001, part 3: p<0.001, HE: part 1: p=0.014, part 2: p=0.006, part 3: p=0.021, CAF: part 1: p=0.008, part 2: p=0.041, Post: part 1: CAF vs CTR: p=0.011, HE vs CTR: p=0.040, part 2: CAF vs CTR: p=0.004, part 3: CAF vs CTR: p=0.022).

Analysis of neural activity during appetitive behaviors such as sniffing revealed that CTR- and HE-rats demonstrated an elevation in ΔF/F at the onset of a sniffing episode, whereas CAF-exposed rats showed attenuated activity (Fig S2C). This resulted in a lack of effect on the AUC post versus pre sniffing (Fig S2D, AUC: Diet: F_(2,34)_=0.717, p=0.496, Epoch: F_(1,34)_= 13.456, p<0.001, Diet*Epoch: F_(2, 34)_=0.706, p=0.500, post-hoc: Pre vs Post: CTR: p=0.012, HE: p=0.042).

Our findings suggest that prolonged exposure to HE and CAF diets significantly impairs the VTA’s response to food rewards and predictive cues associated with food. Typically, an elevation in VTA activity reflects the perceived value of a cue associated with a reward. However, this response is compromised in rats consuming HE or CAF diets, suggesting that the value of food rewards might be diminished in these groups. Moreover, long-term CAF diet intake diminishes VTA responsiveness to both the anticipation and consumption of food rewards more profoundly compared to exposure to a HE diet. This underscores that the *choice of palatable food*, in combination with fat and sugar content (see Table 1, S1, S2, and S3), is a major contributor to disrupting the neural mechanisms governing rewarding behaviors.

### 2.3 Long-term CAF diet also affects VTA neural responses to other natural rewards

We next investigated whether long-term CAF diet consumption influences both neural and behavioral responses to a natural reward other than food. Therefore, we conducted three additional behavioral reward tests using a sexual partner (a sexually experienced male rat) as the reward. Due to the potential occurrence of pseudopregnancies in naturally cycling females, which prevents copulation for approximately two weeks, all three sexual reward tests were conducted on a single day. Unfortunately, not all females reached the estrus phase, and these females were excluded from the analysis. Only six out of ten females in the HE group reached estrus, with two of these females not engaging in copulation. Given the insufficient copulation occurrences in the HE group and the minimal differences from CTR rats in food reward responses, we focused our analysis on CTR and CAF rats. In the CAF group, ten females reached estrus, but three did not exhibit paracopulatory behaviors during the first sexual reward test, suggesting that CAF diet consumption may delay the onset of the receptive phase. To ensure proper inter-rat comparisons, we analyzed photometry and behavioral data from the *best sexual reward* test, defined as the test with the highest number of paracopulatory behaviors.

First, we examined the VTA response to the light cue that preceded door opening. As observed in the food reward test, the light cue elicited a more pronounced increase in neural activity in CTR rats compared to CAF rats (Fig 3A). While the AUC increased significantly after the light cue in CTR rats , no such effect was found in CAF rats (Fig 3B, Light: Diet: F_(2,24)_=2.085, p=0.162, Epoch: F_(1,24)_=14.409, p<0.001, Diet*Epoch: F_(2, 24)_=2.597, p=0.120, post-hoc: Pre-light vs Light: CTR: p=0.002, Light: CAF vs CTR: p=0.041, Door: Diet: F_(2,24)_=3.307, p=0.081, Epoch: F_(1,24)_=2.454, p=0.130, Diet*Epoch: F_(2, 24)_=0.074, p=0.788)).

We also explored activity patterns of the VTA during sexual behaviors. Female rats exhibit paracopulatory behaviors, including hops and darts (Heijkoop, Huijgens, & Snoeren, 2018; Snoeren, Veening, Olivier, & Oosting, 2014; Snoeren, 2019). When aligning the start of a dart (as the moment the female started running before ending in the typical dart position) to the GCaMP6s signaling, we found an increase in VTA activity upon dart initiation in CTR rats, whereas CAF rats showed an attenuated VTA response (Fig 3C/D, AUC: Diet: F_(1,22)_=2.310, p=0.143, Epoch: F_(1,22)_=15.669, p<0.001, Diet*Epoch: F_(1,22)_=2.107, p=0.161, post-hoc: Pre vs Post: CTR: p=0.001, CAF vs CTR: Post: p=0.047). No such effects were observed for short hops (Fig 3E, AUC: Diet: F_(1,20)_=3.201, p=0.089, Epoch: F_(1,20)_=0.246, p=0.625, Diet*Epoch: F_(1,20)_=0.392, p=0.539).

Behaviorally, however, no differences were observed during the sexual reward test. CAF rats exhibited a comparable number of darts (Fig 3G, *t*=0.282, p=0.781) and hops (Fig 3H, *t*=1.088, p=0.291) as CTR rats. Moreover, no differences were found in the number of sexual bouts (Fig 3F and 3I, *t*=0.733, p=0.473), mean duration of sexual bouts (Fig 3J, *t*=0.576, p=0.572), or mean duration of time-outs (Fig 3K, *t*=0.392, p=0.700). Consequently, CAF females received a similar number of copulations as CTR females (Fig 3KL, *t*=-0.371, p=0.715).

While paracopulatory behavior (darts and hops) is often defined as appetitive or proceptive behavior, lordosis is considered a consummatory or receptive behavior. Our data revealed that long-term CAF diet exposure did not alter lordosis quotient (Fig 4A, *t*=-0.425, p=0.676) and lordosis score (Fig 4B, *t*=0.778, p=0.447) in CTR and CAF rats. Similarly, no change in VTA activity was observed during lordosis behavior (Fig 4C). However, when classifying the lordosis behaviors based on intensity (Hardy & Debold, 1971) and type of stimulation, we found that high intensity lordoses resulted in VTA activity increases (Fig 4D). Since the response was limited to the two seconds after the end of a lordosis, we analyzed the AUC for the 2 seconds pre and post lordosis. Unfortunately, no significant effects (Pre vs Post: CTR: p=0.059) were detected in the AUC due to substantial variability (Fig 4D, AUC: Diet: F_(1,12)_=0.651, p=0.435, Epoch: F_(1,12)_=2.265, p=0.158, Diet*Epoch: F_(1,12)_=2.078, p=0.175). Low intensity lordosis responses, on the other hand, failed to elicit changes in VTA activity (Fig 4E, AUC: Diet: F_(1,18)_=0.076, p=0.786, Epoch: F_(1,18)_=0.011, p=0.916, Diet*Epoch: F_(1,18)_=0.091, p=0.767).

**Figure 4.**
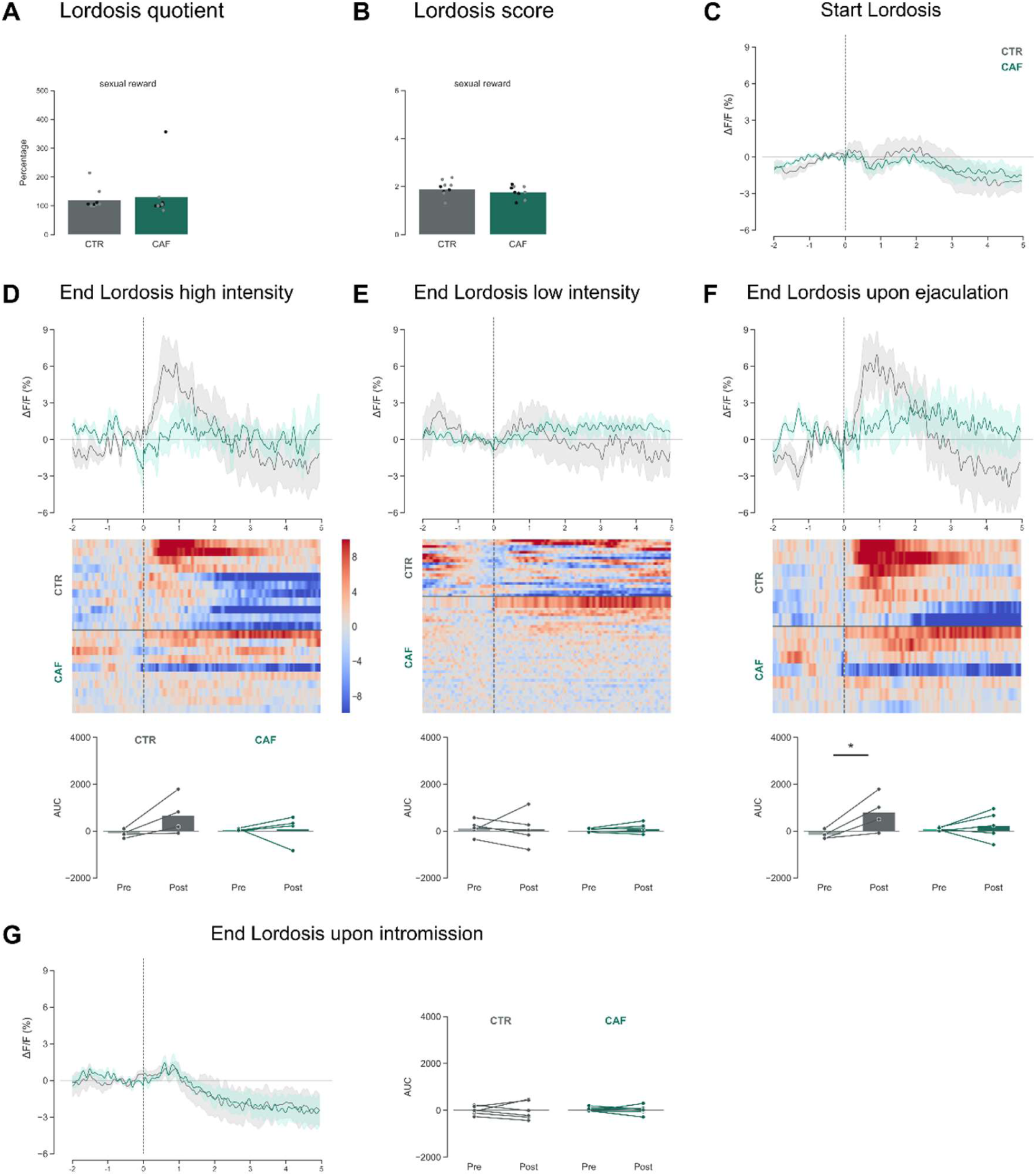
The effects of Cafeteria diet on the ventral tegmental area and lordosis responses during a sexual reward test. **A)** Lordosis quotient. **B)** Lordosis score. **C)** ΔF/F aligned to the start of lordosis behavior. **D)** Upper panel: ΔF/F aligned to the end of high-intensity lordosis behavior. Middle panel: heat maps of ΔF/F aligned to the high-intensity lordosis behavior. Lower panel: AUC pre and post the end of high-intensity lordosis behaviors. **E)** Upper panel: ΔF/F aligned to the end of low-intensity lordosis behavior. Middle panel: heat maps of ΔF/F aligned to the low-intensity lordosis behavior. Lower panel: AUC pre and post the end of low-intensity lordosis behavior. **F)** Upper panel: ΔF/F aligned to the end of lordosis behaviors upon ejaculation. Middle panel: heat maps of ΔF/F aligned to the lordosis behaviors upon ejaculations. Lower panel: AUC pre and post the end of lordosis behaviors upon ejaculation. **G)** Left panel: ΔF/F aligned to the end of lordosis behaviors upon intromission. Right panel: AUC pre and post the end of lordosis behaviors upon intromission. **All** Figures show rats on control (CTR, grey color bars and lines) and cafeteria (CAF, green color bars and lines) diets during the reward phase of the best sexual reward test. The dots in the behavioral data have colors representing their body weight category (grey: lighter than average, black: heavier than average). The lines in the ΔF/F figures represent the mean and shaded areas the SEM. CTR n=6, CAF n=8. * p<0.05 compared to by line connected group. ^a^ p<0.05 compared to same bar of CTR.

When we analyzed the photometry data based on copulation type, we found a VTA activity increase after receiving an ejaculation in CTR females (Fig 4F), resulting in a significant change in the AUC from before and after the lordosis response in CTR rats (Fig 4F, AUC: Diet: F_(1,14)_=0.442, p=0.517, Epoch: F_(1,14)_=5.622, p=0.033, Diet*Epoch: F_(1,14)_=3.028, p=0.104, post-hoc: Pre vs Post: CTR: p=0.015). CAF rats did not show this activity increase, once again demonstrating an attenuated VTA response. No activity changes were observed following a lordosis response to intromission (Fig 4G, Diet: F_(1,22)_=0.175, p=0.680, Epoch: F_(1,22)_=0.039, p=0.845, Diet*Epoch: F_(1,22)_=0.026, p=0.874). Overall, this suggests that the VTA does not encode the onset of lordosis behavior, but rather responds to the received stimulation.

In summary, our findings indicate a role for the VTA in responses to both food and sexual rewards. While long-term exposure to junk food profoundly affected behavior to food rewards, normal levels of behavior were observed for sexual rewards. Still, rats exposed to CAF diet showed attenuated VTA activity in response to both food and sexual rewards.

### 2.4 Two weeks of standard chow diet is insufficient to reverse the CAF-induced changes in response to food rewards

To assess the reversibility of neural and behavioral alterations induced by long-term exposure to CAF diet, all rats were transitioned to a standard chow diet for two weeks following their last behavioral test. This was followed by three iterations of the behavioral reward test with a food stimulus (CAF) and one instance with a sexual stimulus (Fig 5A). We observed that the two-week period of dietary reversal was insufficient to restore the attenuated VTA responses elicited by the light cue associated with a food reward in CAF-exposed rats (Fig 5B/C, AUC: Light: Diet: F_(2,26)_=4.273, p=0.049, Epoch: F_(1,26)_=27.659, p<0.001, Diet*Epoch: F_(2, 26)_=5.794, p=0.023, post-hoc: Pre-light vs Light: CTR: p<0.001, CAF: p=0.033, Light: CAF vs CTR: p=0.004, Door: Diet: F_(2,26)_=3.223, p=0.084, Epoch: F_(1,26)_=7.192, p=0.013, Diet*Epoch: F_(2, 26)_=0.839, p=0.368, post-hoc: Pre-light vs Light: CTR: p=0.028).

**Figure 5.**
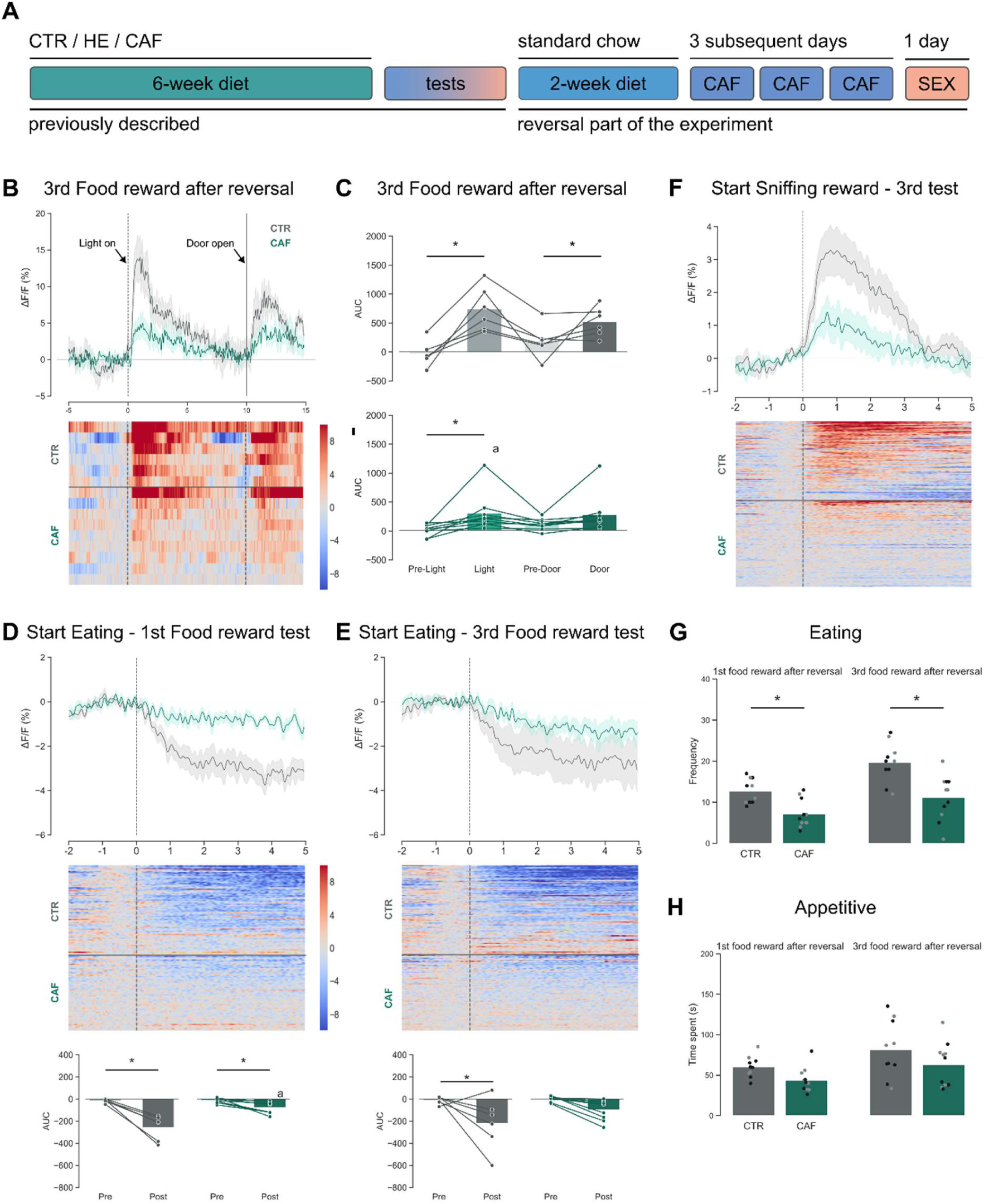
The effects of reversal diet after Cafeteria diet exposure on the ventral tegmental area and behavioral responses during a food reward test **A)** Schematic representation of the experimental design. **B)** Upper panel: ΔF/F aligned to the light cue at t=0 and door opening at t=10s. Lower panel: heat maps of ΔF/F aligned to the light cue and door opening. **C)** AUC before (pre-Light) and after (Light) the light cue, and before (Pre-door) and after (Door) the door opening. **D)** Upper panel: ΔF/F aligned to the start of eating. Middle panel: heat maps of ΔF/F aligned to the start of eating. Lower panel: AUC pre and post the start of eating. **E)** Upper panel: ΔF/F aligned to the start of eating. Middle panel: heat maps of ΔF/F aligned to the start of eating. Lower panel: AUC pre and post the start of eating. **F)** Upper panel: ΔF/F aligned to the start of sniffing the reward. Lower panel: heat maps of ΔF/F aligned to the start of sniffing. **G)** Total number of eating episodes. **H)** Time spent on appetitive behaviors. **All** Figures show rats on control (CTR, grey color bars and lines) and cafeteria (CAF, green color bars and lines) diets during the reward phase of the food reward tests after reversal diet. The dots in the behavioral data have colors representing their body weight category (grey: lighter than average, black: heavier than average). The lines in the ΔF/F figures represent the mean and shaded areas the SEM. CTR n=6, CAF n=9 for fiber photometry data and CTR n= 10, CAF n=11 for behavioral data. The dots in the behavioral data have colors representing their body weight category (grey: lighter than average, black: heavier than average) * p<0.05 compared to by line connected group. ^a^ p<0.05 compared to same bar of CTR.

A similar lack of recovery was evident in the VTA neural responses during the behavioral interactions with the food reward. Specifically, CAF-exposed rats continued to exhibit attenuated neural activity responses during eating in both the 1^st^ (Fig 5D) and 3^rd^ food reward test (Fig 5E) post-reversal diet. The attenuated response persisted in CAF rats during the initial food reward test after dietary reversal, as demonstrated by a significant difference in AUC (Fig 5D, AUC: Diet: F_(1,26)_=17.511, p<0.001, Epoch: F_(1,26)_=46.771, p<0.001, Diet*Epoch: F_(1,26)_=15.952, p<0.001, post-hoc: Pre vs Post: CTR: p<0.001, CAF: p=0.033, Post: CAF vs CTR: p<0.001), but this effect diminished by the third food reward test (Fig 5E, AUC: Diet: F_(1,26)_=2.484, p=0.127, Epoch: F_(1,26)_=12.877, p=0.001, Diet*Epoch: F_(1,26)_=1.678, p=0.207, post-hoc: Pre vs Post: CTR: p=0.004), even though it remained visible in the heatmaps. These results suggest a persistent effect of long-term CAF diet on the VTA, which a two-week reversal diet cannot readily undo.

Interestingly, even though two weeks of healthy diet was not sufficient to restore VTA function, a partial behavioral recovery was observed. Analysis revealed that CAF rats still ate less frequently (Fig 5G, Diet: F_(2,19)_=18.799, p<0.001, Test: F_(1,19)_=35.494, p<0.001, Diet*Test: F_(1,19)_=2.442, p=0.135, post-hoc: CAF vs CTR: 1^st^ test: p=0.006, 3^rd^ test: p<0.001, 1^st^ vs 3^rd^ test: CTR: p<0.001, CAF: p=0.005) and had fewer eating bouts (Fig S3A, Diet: F_(2,19)_=7.586, p=0.013, Test: F_(1,19)_=6.038, p=0.024, Diet*Test: F_(1,19)_=0.014, p=0.908, post-hoc: CAF vs CTR: 1^st^ test: p=0.023, 3^rd^ test: p=0.018) than CTR rats in both the 1^st^ and 3^rd^ food reward test following the dietary reversal period. However, regarding time spent eating (Fig S3B, Diet: F_(2,19)_=6.640, p=0.018, Test: F_(1,19)_=26.290 p<0.001, Diet*Test: F_(1,19)_=0.303, p=0.588, post-hoc: CAF vs CTR: 1^st^ test: p=0.021, 1^st^ vs 3^rd^ test: CTR: p=0.005, CAF: p<0.001), or mean duration of time-outs (Fig S3C, Diet: F_(2,17.9)_=6.009, p=0.025, Test: F_(1,17.309)_=6.282 p=0.022, Diet*Test: F_(1,17.309)_=0.578, p=0.457, post-hoc: CAF vs CTR: 1^st^ test: p=0.017, 1^st^ vs 3^rd^ test: CAF: p=0.033), CAF rats displayed diminished levels compared to CTR rats only in the 1^st^ food reward test, with no significant differences observed in the third test. Furthermore, consistent with pre-reversal observations, all rats spent an equal amount of time within an eating bout in all tests (mean duration of eating bouts, Fig S3D, Diet: F_(2,19)_=0.201, p=0.659, Test: F_(1,19)_=19.547 p<0.001, Diet*Test: F_(1,19)_=0.832, p=0.373, post-hoc: 1^st^ vs 3^rd^ test: CTR: p=0.025, CAF: p=0.001). Likewise, although behavioral indications suggested a recovery in appetitive behavior (approaching and sniffing the food reward) among CAF rats in the 1^st^ and 3^rd^ food reward test post-reversal (Fig 5H, Diet: F_(2,19)_=4.016, p=0.060, Test: F_(1,19)_=11.974, p=0.003, Diet*Test: F_(1,19)_=0.008, p=0.931, post-hoc: 1^st^ vs 3^rd^ test: CTR: p=0.024, CAF: p=0.024), neural responses within the VTA remained attenuated compared to CTR rats at both the 1^st^ (Fig 5E, AUC: Diet: F_(1,26)_=17.511, p<0.001, Epoch: F_(1,26)_=46.771, p<0.001, Diet*Epoch: F_(1,26)_=15.952, p<0.001, post-hoc: Pre vs Post: CTR: p<0.001, CAF: p=0.033, Post: CAF vs CTR: p<0.001) and 3^rd^ food reward test (Fig 5F, AUC: Diet: F_(1,26)_=2.484, p=0.127, Epoch: F_(1,26)_=12.877, p=0.001, Diet*Epoch: F_(1,26)_=1.678, p=0.207, post-hoc: Pre vs Post: CTR: p=0.004).

Altogether, our findings suggest that a two-week period of healthy dietary intervention is insufficient to fully restore the immediate behavioral or neural responses to a CAF food reward. However, it does partially improve behavioral effects after repeated exposures, *despite* persistent reduced responsiveness in the VTA.

### 3.5 Two weeks of standard chow diet inadequate in restoring the CAF-induced changes to sexual rewards

Lastly, following a two-week exposure to a healthy diet, the rats underwent another sexual reward test. Once more, CAF-exposed rats exhibited no recovery in VTA responsiveness to the light-cue, maintaining an attenuated response compared to control rats (Fig S4A/B, AUC: Light-cue: Diet: F_(1,22)_=1.863, p=0.186, Epoch: F_(1,22)_=11.247, p=0.003, Diet*Epoch: F_(1, 22)_=1.787, p=0.195, post-hoc: Pre-light vs Light: CTR: p=0.010, Door: Diet: F_(1,22)_=0.013, p=0.911, Epoch: F_(1,22)_=3.480, p=0.075, Diet*Epoch: F_(1,22)_=0.007, p=0.933).

Furthermore, the two weeks of healthy diet did not restore the VTA responsiveness in relation to sexual behavior. While there was still no difference in sexual behavior between CTR and CAF rats (Fig S4C-E, Darts: *t*=-1.131, p=0.273, Hops: *t*=-0.161, p=0.874 (data not shown), Lordosis quotient: *t*=-1.232, p=0.236, Lordosis score: *t*=-1.197, p=0.249), CAF rats still showed an attenuated VTA response upon darting behavior compared to CTR rats after reversal of diets (Fig S4F, AUC: Darting: Diet: F_(1,20)_=7.066, p=0.015, Epoch: F_(1,20)_=23.791, p<0.001, Diet*Epoch: F_(1,20)_=4.414, p=0.049, post-hoc: Pre vs Post: CTR: p<0.001, CAF: p=0.026, CAF vs CTR: Post: p=0.003, Hopping (data not shown): Diet: F_(1,18)_=0.215, p=0.649, Epoch: F_(1,18)_=6.672, p=0.019, Diet*Epoch: F_(1,18)_=1.274, p=0.274, post-hoc: Pre vs Post: CTR: p=0.032,).

Moreover, the data revealed that CAF rats continued to exhibit a reduced neural response in the VTA during lordosis episodes. While there was an insufficient trial number of the highest lordosis intensity in CTR rats for a robust comparison, the second-highest intensity (Fig S4G) and now also the low intensity of lordosis (Fig S4H) still evoked an increase in activity in CTR rats immediately following the end of a lordosis. However, these increases were too subtle to elicit significant effects in the AUC (data not shown). Likewise, the dietary reversal failed to restore the attenuated responses of CAF rats regarding received ejaculations (Fig S4I).

## 4. Discussion

Our study reveals that rats on a long-term CAF diet, which consists of a *free choice* of highly palatable and unhealthy products, exhibited reduced appetitive and consummatory behavior towards the CAF reward compared to rats on a control diet. These behavioral changes were accompanied by diminished neural activity in the VTA in response to a food-predictive cue and when the rats interacted with the food reward directly. Similarly, reductions in VTA activity were observed in relation to a sexual reward. However, no significant behavioral differences were observed between the control and CAF rats in terms of appetitive (darting and hopping) or consummatory (lordosis) behaviors during sexual interactions. Furthermore, the study supported our hypothesis that CAF diet alters the functioning of the reward circuitry more severely than a high energy diet. This finding coincided with the marked difference that CAF and HE diets had on food reward-related behavior: while long-term exposure to CAF limited interactions with the food reward, exposure to HE did not. Finally, we found that a reversal diet intervention of two weeks with healthy food was not sufficient to restore the behavioral and neural alterations seen after CAF diet consumption.

Overall, these results suggest that prolonged exposure to junk food leads to diminished activity of VTA neurons upon both food and sexual rewards, which corresponds with behavioral changes in response to food rewards but not sexual rewards.

Our data support our hypothesis that CAF diet more severely affects the VTA than HE diet does. This effect is most likely due to the element of *choice* of unhealthy products with different taste, texture, novelty and variety in CAF diet in contrast to HE pellets (Lalanza & Snoeren, 2021; Sampey, et al., 2011). Consistent with other studies, we observed that the free choice consumption and variation of food items led to greater body weight gain (See Table 3) (la Fleur, et al., 2014) and increased overeating (Rolls, et al., 1981) (See Table S2). Notably, each cage showed different consumption patterns, with preferences for different foods (shown in Table S3). This overeating may contribute to more pronounced alterations in the brain, as indicated by stronger attenuation of VTA responses in CAF-fed rats compared to those on the HE diet. Thus, CAF diet seems to be a more robust model to study excessive junk food consumption than HE pellets, even though implementing it is more laborious and provides less control over the nutritional value of the consumed food. Future experiments could conduct a more detailed analysis of the macronutrient mix consumed by individual rats to provide insights into how specific dietary components influence VTA responsiveness and behavior.

To our knowledge, this is the first study showing that long-term CAF diet can change event-related neural responses in the VTA toward a food and sexual reward, as well as reward-predictive cues. So far, most studies have shown effects of junk food on the hippocampus or nucleus accumbens, but only a few studies have reported potential changes in the VTA (Bourdy, et al., 2021; Carlin, et al., 2016; Martire, et al., 2014; Sharma, Fernandes, & Fulton, 2013). Interestingly, all these studies report reduced levels of receptors (Bourdy, et al., 2021; Carlin, Hill-Smith, Lucki, & Reyes, 2013; Martire, et al., 2014) or tyrosine hydroxylase (Sharma, et al., 2013), supporting our findings of reduced VTA functioning after junk food exposure. In addition, it was recently found that the ways dopamine neurons in the VTA encode food and opposite-sex social motivation show overlap (Willmore, et al., 2023). This study supports our hypothesis that desensitization of the VTA by excessive consumption of CAF diet can disrupt the neural responses toward other natural rewards as well, as shown in our study.

Our findings are in support of the reward hyporesponsivity theory that long-term exposure to foods rich in fat and sugar can cause a blunted behavioral reward response (Ducrocq, et al., 2019; Johnson & Kenny, 2010; Stice, Spoor, Bohon, & Small, 2008). While some studies have shown an increase in motivation to work for a food reward (Greenwood, et al., 1974; la Fleur, et al., 2007;

Narayanaswami, et al., 2013; Robinson, et al., 2015; Vasselli, et al., 1980), or no effect at all (Spaulding, et al., 2024), more studies found that rats on a junk food diet show reduced interest in cues for a food reward and show attenuated operant responses (Blaisdell, et al., 2014; Davis, et al., 2008; Ducrocq, et al., 2019; Harb & Almeida, 2014; Ibias, et al., 2016; Rossi, et al., 2013; Steele, et al., 2019; Tantot, et al., 2017; Tracy, Wee, Hazeltine, & Carter, 2015), or longer latencies to collect a reward (Harb & Almeida, 2014; Shin, et al., 2011). In contrast to most of these studies, our experimental design permitted free access to, and interaction with the reward and is therefore considered a ‘low effort’ paradigm. In this regard, our findings are in line with the other studies in which little or no effort was required to obtain the rewards (Duca, et al., 2014; Ducrocq, et al., 2019; Johnson, 2012; la Fleur, et al., 2007; Vucetic, et al., 2011).

Since our behavioral reward test included a light cue indicating free access to the reward, our set-up enabled us to study the effects of excessive junk food consumption on the VTA both during classical conditioning and during reward approach and consumption. Initially, in the first test in all groups, the VTA responded to the light cue in a way previously described for novel and neutral cues (Horvitz, Stewart, & Jacobs, 1997). By the fifth and final test however, VTA activity in the control group was higher in reaction to the cue light, relative to the first test, suggesting that the cue light had become a conditioned stimulus. In contrast, light-evoked activity in the VTA for both the CAF and HE groups did not differ from the first food reward test. Furthermore, all groups had elevated activity levels following opening of the door, which provided access to the reward, during the fifth food reward test, but not the first. We interpreted this response as a part of the cue-reward sequence that was not yet fully predicted by the cue light, and this might suggest that the VTA is mainly affected in its role to encode the predictive and incentive properties of a reward rather than in its role to signal novel stimuli that could lead to potential rewards (Day, Roitman, Wightman, & Carelli, 2007; Horvitz, et al., 1997; Ljungberg, Apicella, & Schultz, 1992). After experiencing decreased pleasure and desire for the reward, the adjusted reward prediction after Pavlovian conditioning should typically lead to diminished VTA responses to the light-cue and reward (Schultz, 2015, 2016; Schultz, Dayan, & Montague, 1997). We hypothesize that this is what happens in the rats exposed to CAF and HE diets.

The lack of behavioral changes in responses to a sexual reward, on the other hand, was surprising. The observation that VTA activity was attenuated despite the absence of explicit changes in behavioral responses to a sexual reward, does support the notion that while long-term CAF exposure may exert effects on some of the same anatomical substrates of different natural rewarding behaviors, redundancy by means of other pathways might contribute to the robustness of sexual behavior. We have previously shown how different natural rewards can recruit similar brain networks to regulate the display of motivated behaviors (Huijgens, et al., 2024). Still, the actual display of either feeding or sexual behavior is also regulated by two different and specialized circuitries. While the lateral hypothalamus plays a crucial role in feeding behavior (Anand & Brobeck, 1951; Stuber & Wise, 2016), the ventromedial nucleus of the hypothalamus is strongly implicated in regulating female sexual behavior (Matthews, et al., 1997; Pfaff & Sakuma, 1979; Snoeren, et al., 2015; Snoeren, 2019; Yin & Lin, 2023). With dense projections between the VTA and the feeding and sexual behavior brain networks (Swanson, 1982), it is possible that CAF diet exposure also affects other regions than the VTA related to feeding behavior, while having less effect on the sexual behavior related regions. Alternatively, changes in sexual motivation might be found using more subtle behavioral tests, in which consummatory behaviors are not displayed, such as the sexual incentive motivation test (Heijkoop, et al., 2018; Huijgens, Heijkoop, & Snoeren, 2023), or a test in which the rats have to work to gain access to the sexual reward. It is also plausible that prolonged testing durations can unveil behavioral disparities between CTR and CAF rats. For example, if the motivation to continue copulation is affected, it would likely become more apparent with longer test durations. Another explanation for the lack of effects on sexual behavior can be the fact that our female rats were sexually experienced at the start of testing. Beloate et al. (2016) have shown that inhibition of dopaminergic neurons in the VTA does not change the initiation and performance of sexual behavior in male rats, but that it does prevent sexual experience to affect neuronal plasticity and cross-sensitization of amphetamine-induced conditioned place preference (Beloate, et al., 2016). Following this line of thought, it might be possible that the diminished CAF-induced VTA responses might lead to altered sexual behavior in sexually naïve female rats as well. Furthermore, it would be interesting to explore the effects on male rat sexual behavior.

Furthermore, we would like to stress that to our knowledge, this is the first study to investigate how the VTA is activated during female sexual behavior. In our study, we found that an increase in neural activity in the VTA was linked to the start of darting, but not hopping events. The increase in VTA activity was aligned to the the initiation of running for the darting response, rather than the moment the rats positioned themselves on the ground for the male to mount. Regarding lordosis behavior, on the other hand, VTA activity was aligned to the end of a lordosis behavior. Lordosis behavior is a response toward a copulatory behavior by the male and when this behavior has ended, an increase in VTA activity was seen exclusively after an ejaculation. This suggests that the VTA has a stimulation-specific response and that it might encode the rewarding elements perceived from such a stimulation.

Finally, we found that a 2-weeks reversal to a healthy diet was insufficient to restore the behavioral and neural responses caused by long-term CAF diet. This finding is in line with another study that reported that two weeks of diet withdrawal did not restore the behavioral changes caused by CAF diet exposure (Johnson & Kenny, 2010). Interestingly though, other studies have reported binge-like eating behavior after withdrawal from junk food diets (Carlin, et al., 2016; Sharma, et al., 2013).

Others found a restoration of sucrose preference after 4 weeks of chow feeding (Carlin, et al., 2016). More research is needed to determine whether the behavioral changes will return to normal after longer exposure to healthy chow diet. The findings of unchanged, blunted VTA responses, on the other hand, seem to be rather consistent with other studies that evaluated the persistence of neural changes caused by junk food exposure. It was found that D1 receptor expression and TH levels remained lower after abstinence from CAF diet (Ong, et al., 2013; Sharma, et al., 2013). Interestingly though, a sex difference was present in these findings. While female rats maintained reduced D1 receptor expression levels, males seem to return to baseline after 72 hours (Ong, et al., 2013).

One limitation of our study was the use of a non-specific GCaMP promotor (hSyn) for the fiber photometry recordings. At the time the study was conducted, no successful cell-specific promotors were available for rats. While transgenic mice are often used to target specific neuronal populations, the behavioral profile in rats offers advantages in this type of study. In addition, rats are a better translational model in studies of obesity and effects of dietary manipulations (Buettner, Scholmerich, & Bollheimer, 2007). Although cell specificity was not reached in this study, previous studies have shown that the VTA consists of both dopaminergic, glutamatergic and GABAergic cells, and many neurons co-release different neurotransmitters (Conrad, Oriol, Kollman, Faget, & Hnasko, 2024; Dobi, Margolis, Wang, Harvey, & Morales, 2010; Ma, Zhong, Liu, & Wang, 2023; Morales & Margolis, 2017; Nair-Roberts, et al., 2008). It is therefore impossible to determine which population is of most importance for our findings of an attenuated VTA response after prolonged junk food consumption. However, since most neurons are dopaminergic (Nair-Roberts, et al., 2008) and our responses to cues and rewards are consistent with responses seen when people have measured dopamine neuron firing (Schultz, et al., 1997) or dopamine release with voltammetry (Roitman, et al., 2008), it is likely that dopaminergic neurons play an important role in our findings. Future studies should reveal the role of the different neurons in the VTA in more detail.

In conclusion, the results indicate that long-term junk food exposure severely affects the reward system by altering behavioral responses to a CAF food reward and desensitizing the VTA to a food and sexual reward. Exposure to a CAF diet in which rats are exposed to a choice of foods rich in fat and sugar has more severe consequences than a HE diet. Finally, two weeks of healthy diet is not sufficient to restore the behavioral and neural changes. Our findings support the notion that excessive junk food consumption can have undesirable long-lasting consequences.

## Acknowledgements

Financial support was received from Helse Nord (HNF1443-19) and MSCA-IF (882946). We thank Carina Sørensen, Lorenzo Ragazzi, Ragnhild Osnes, Remi Osnes and Siri Knudsen for the excellent care of the animals, and Truls Traasdahl and Thomas Nermo for the construction of the behavioral reward test box. Finally, we thank the Advanced Microscopy Core Facility of UiT The Arctic University of Tromsø for access to their equipment for the brain analyses.

## Author contributions

JFL: Experimental design, data gathering, behavioral annotation, data curation, analysis, writing – review and editing, funding acquisition

JCO: Data gathering, behavioral annotation, analysis, writing – review and editing

PTH: Methodology, writing – review and editing

JEM: Methodology, writing - review and editing

RH: Experimental design, methodology, data gathering, data curation, analysis, supervision, writing – original draft, funding acquisition

EMSS: Experimental design, methodology, data gathering , data curation, Programming/software, analysis, supervision, writing – original draft, funding acquisition

