## Supplementary information for "Behavioral and neural alterations of the ventral tegmental area by exposure to junk food in rats"

### Supplementary methods

**Sexual Experience.**

The natural estrous cycle of the females was monitored using an impedance meter (Muromachi MK-12, Japan)(Koto, et al., 1987). When in estrus, the females obtained sexual experience in a copulation box (60 x 40 x 40 cm, with a Plexiglas front) in a room with dimmed lighting conditions. A receptive female was first habituated to the copulation box for five minutes. Following this habituation, a male was introduced, and the session continued until ejaculation was achieved or for a maximum of 30 minutes. An ejaculation was characterized by a prolonged intromission accompanied by additional muscle contraction during climax.

**Behavioral reward test box**

The behavioral reward test was conducted in a box (75x30x70 cm, Fig 1B) in which two chambers (30x30x70 cm and 45x30x70 cm) were separated by a wall containing 2 sliding doors. The side of the dividing wall facing the left chamber (starting chamber, not containing the reward) had a light between the two sliding doors. Small apertures in these doors enabled the subject rat to see, hear and smell the reward in the right chamber. The floors were covered with fresh wood chip bedding. The front wall of the box facing the camera was made of transparent methacrylate.

**Fiber photometry recording and processing**

To assess the activity of VTA neurons during the behavioral reward tests, the bulk fluorescence signal generated by GCaMP6s expressing cells was recorded using fiber photometry(Gunaydin, et al., 2014; Lerner, et al., 2015). Signal processing and acquisition hardware (RZ10x; Tucker-Davis Technologies, Florida, USA) was used to control a 465 nm Lux RZ10x integrated LED for fluorophore excitation, and a 405 nm Lux RZ10x integrated LED for isosbestic excitation. The excitation lights were modulated at 210 Hz (465 nm) and 330 Hz (405 nm), filtered and combined by a fluorescence mini cube (Doric Lenses, Canada), and delivered through a 400 µm core, 0.57 NA, low-autofluorescence fiber optic patch cable (Doric Lenses, Canada, MFP_400/430/1100-0.57_5m_FCM-MF2.5_LAF) to an implanted fiber (Doric lenses, Canada, MFC_400/430-0.66_9mm_MF2.5_FLT). LED power was set at a level which resulted in a ~15 and ~30 mV amplitude output signal for 405 and 465 nm demodulated channels, respectively. All signals were acquired using Synapse Essentials software (Tucker-Davis Technologies, Florida, USA). Fluorescence was sampled at 1017 Hz and demodulated by the processor. The start of the behavioral reward test and the moment the light was turned on were time stamped by registering TTL signals generated by the USB-I/O box 12 system (Noldus, The Netherlands). With the help of The Observer XT 12.5 (most behavioral data) and XT 16 (Novel food reward data, Noldus), the behavioral events were time stamped and aligned with the fiber photometry data.

Data were exported to Python from Synapse using a script provided by Tucker-Davis Technology, Florida, USA. Custom Python scripts were used to independently normalize the demodulated 465 nm and 405 nm signals. To calculate ΔF/F, a least-squares linear fit was applied to the 405 nm signal to align it to the 465 nm signal, producing a fitted 405 nm signal that was used to normalize the 465 nm as follows: ΔF/F = (465 nm signal − fitted 405 nm signal)/fitted 405 nm signal(Lerner, et al., 2015). Fractions of the signal recording in which noisy signals were recorded (e.g., when the cable detached or hit the wall or door) were removed from the recordings when an interquartile range of more than 2 was found. We then aligned the ΔF/F to baseline zero by subtracting the average of a ΔF/F signal collected at the start of each of the events. The area under the curve (AUC) was calculated from this corrected signal. To compare the AUC before and after the event, the AUC was corrected per second.

**Brain processing, immunostaining and imaging**

At the end of the experiments, rats were injected intraperitoneally with a lethal dose of pentobarbital (100 mg/kg), and transcardially perfused with 0.01 M phosphate buffered saline (PBS; pH 7.4) followed by 4% formaldehyde in 0.01 M PBS. Brains were removed and post-fixed in 4% formaldehyde for 48 hours. Then, the brains were transferred to 20% sucrose in 0.01 M PBS for one day and placed in 30% sucrose in 0.01 M PBS. When sunken, the brains were snap frozen in isopentane cooled by dry ice and stored at -80 °C until sectioning on a sliding microtome (SM2010R, Leica Biosystems, Germany) into 40 µm coronal sections. These sections were stored in cryoprotectant solution (30% sucrose w/v, 30% ethylene glycol v/v in 0.01 M PBS, pH 7.4) until further use.

To assess the expression of GCaMP in the VTA, immunohistochemistry was performed on selected sections. These were washed in 0.1M Tris-buffered-saline (TBS), blocked for 30 minutes in 0.5% BSA in TBS, and incubated with polyclonal chicken anti-EYFP (1:100 000), Abcam, UK cat. ab13970) antibody solution containing 0.1% Triton-X and 0.1% BSA in TBS on an orbital shaker for 24 hours at room temperature + 24 hours at 4 °C. The sections were then washed four times in TBS and subsequently incubated in biotinylated goat anti-chicken (1:400, Abcam, UK, cat. ab6876) antibody solution containing 0.1 % BSA in TBS for 30 min, followed by incubation with avidin-biotin-peroxidase complex (VECTASTAIN ABC-HRP kit, Vector laboratories, UK, cat. PK-6100, dilution: 1 drop A + 1 drop B in 10 mL TBS) solution for 30 min, and 3,3′-diaminobenzidine solution (DAB substrate kit (HRP), Vector laboratories, UK, cat. SK-4100, dilution: 1 drop R1 (buffer solution) + 2 drops R2 (3,3′-diaminobenzidine solution) + 1 drop R3 (hydrogen peroxide solution) in 5 mL water) for 5-10 min, with TBS washes between all steps. The slides were left until dry, after which the sections were dehydrated with 50-99.8% ethanol baths, cleared with xylene, and coverslipped using Entellan mounting medium (Sigma, St. Louis, USA).

Finally, the location of the fiber and GCaMP expression were visually verified on the sections with a microscope (Primo Star, Carl Zeiss, Germany). Only the rats that had both GCaMP expression and a fiber within the VTA (between AP -4.7 mm and AP -5.2 mm) were included in the fiber photometry data analysis.

### Supplementary figures

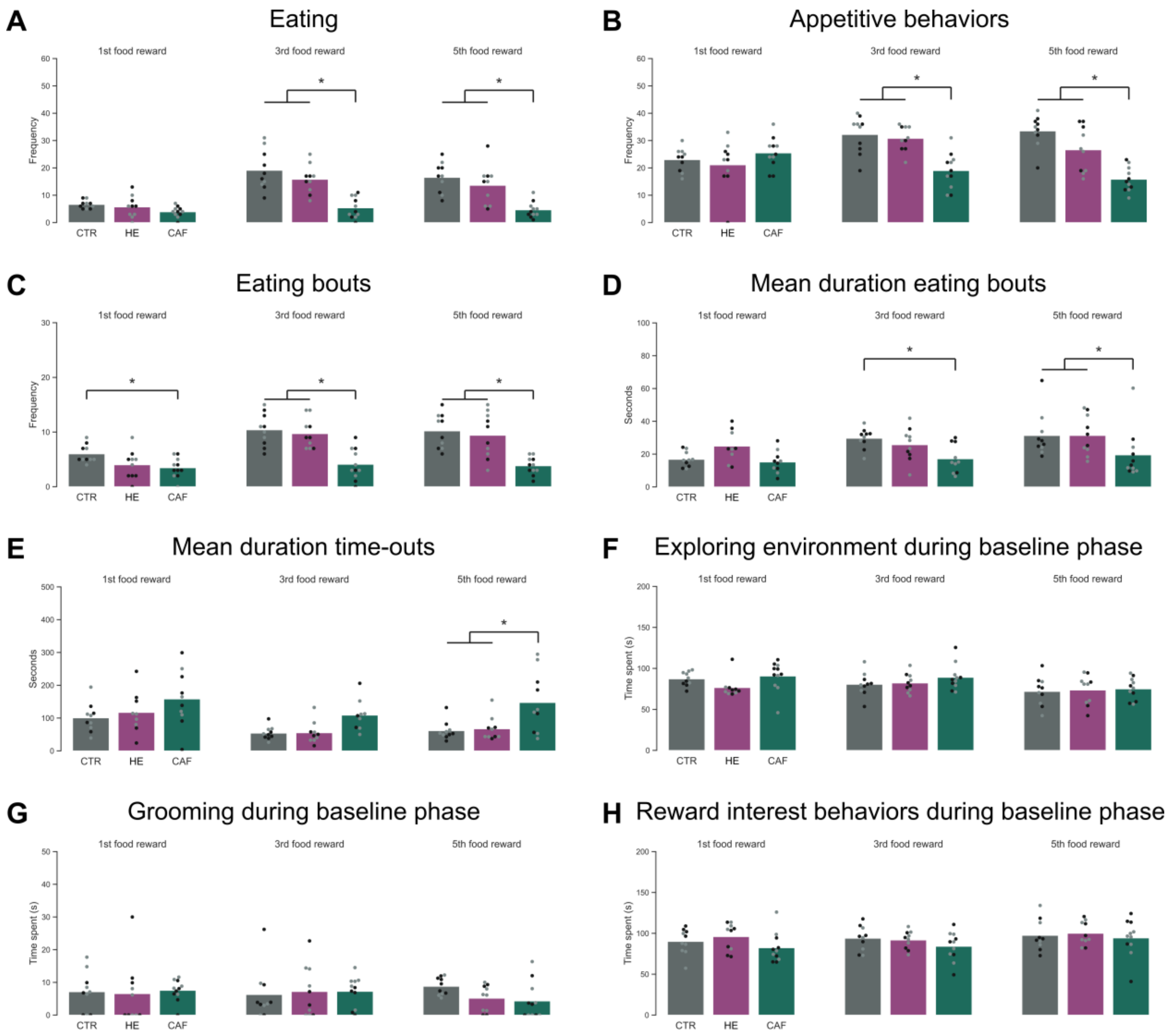


**Figure S1.** The effects of Cafeteria, High energy, and control diet on behavioral responses during a food reward test. **A)** Total number of times eating, **B)** Number of appetitive behaviors, **C)** Total number of eating bouts, **D)** Mean duration of eating bouts, **E)** Mean duration of time-outs, **F)** Time spent exploring the environment, **G)** Time spent grooming, **H)** Time spent on behaviors that indicate interest in the reward in the other chamber. **All** Figures show rats on control (CTR, grey bar), high energy pellet (HE, purple bar), and cafeteria (CAF, green bar) diet, with their dot colors representing their body weight category (grey: lighter than average, black: heavier than average). All figures represent behaviors recorded during the reward phase of the 1$^{st}$, 3$^{rd}$ and 5$^{th}$ behavioral reward test with a food reward, unless stated differently in figure title. CTR n=10, HE n=10, CAF n=11. * $p<0.05$ compared to by line connected group.

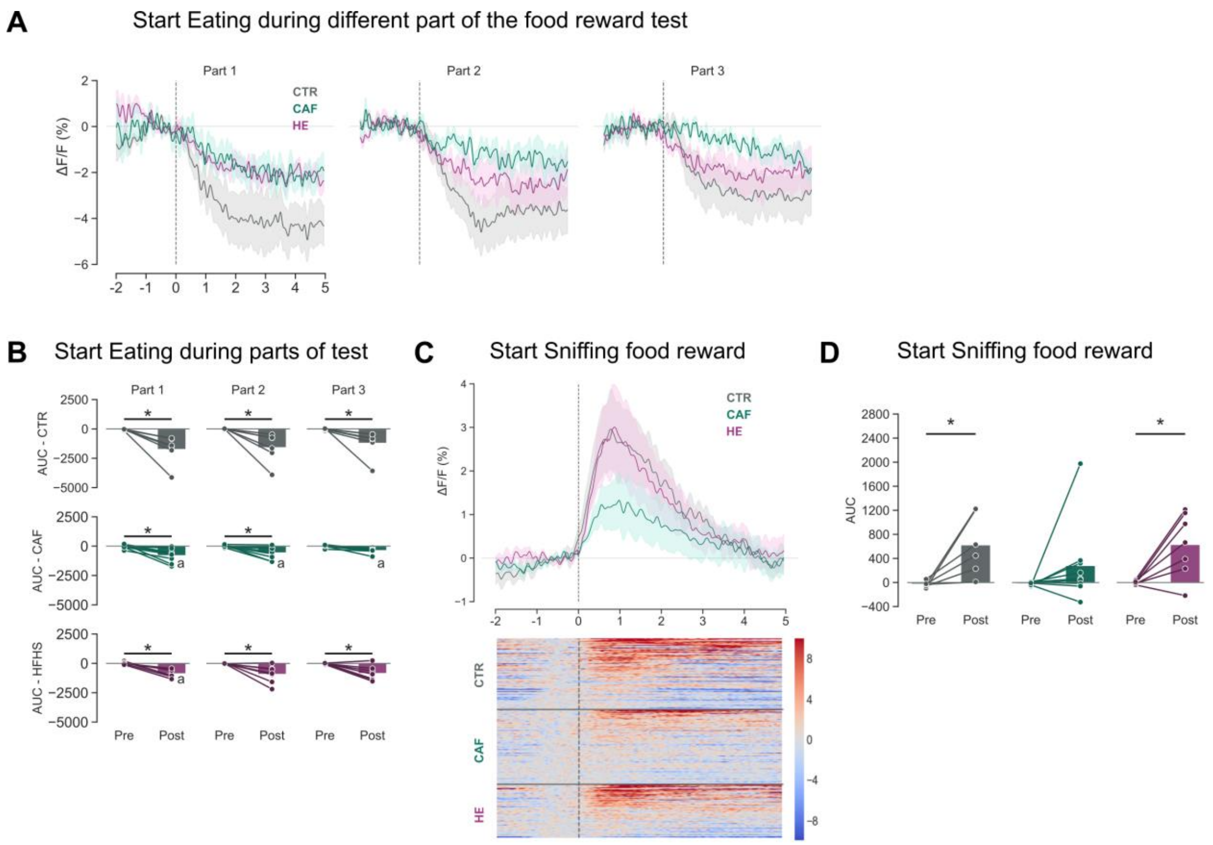


**Figure S2.** The effects of Cafeteria, High energy, and control diet on the ventral tegmental area during a food reward test. **A)** ΔF/F aligned to the start of the start of eating behaviors during the 1st, the 2nd and last part of the reward phase, B**)** AUC pre and post the start of eating behaviors during the 1st, the 2nd and last part of the reward phase. **C)** Upper panel: ΔF/F aligned to the start of sniffing reward. Lower panel: Heat maps of ΔF/F aligned to the start of sniffing reward. Trials are sorted by largest responses, **D)** AUC pre and post the start of sniffing reward behaviors. **All** Figures show rats on control (CTR, grey color bars and lines), cafeteria (CAF, green color bars and lines), and high energy pellet (HE, purple color bars and lines) diet during all food reward tests. The lines in the ΔF/F figures represent the mean and shaded areas the SEM. CTR n=6, CAF n=9, HE n=5. * $p<0.05$ compared to by line connected group. [a] $p<0.05$ compared to same bar of CTR.

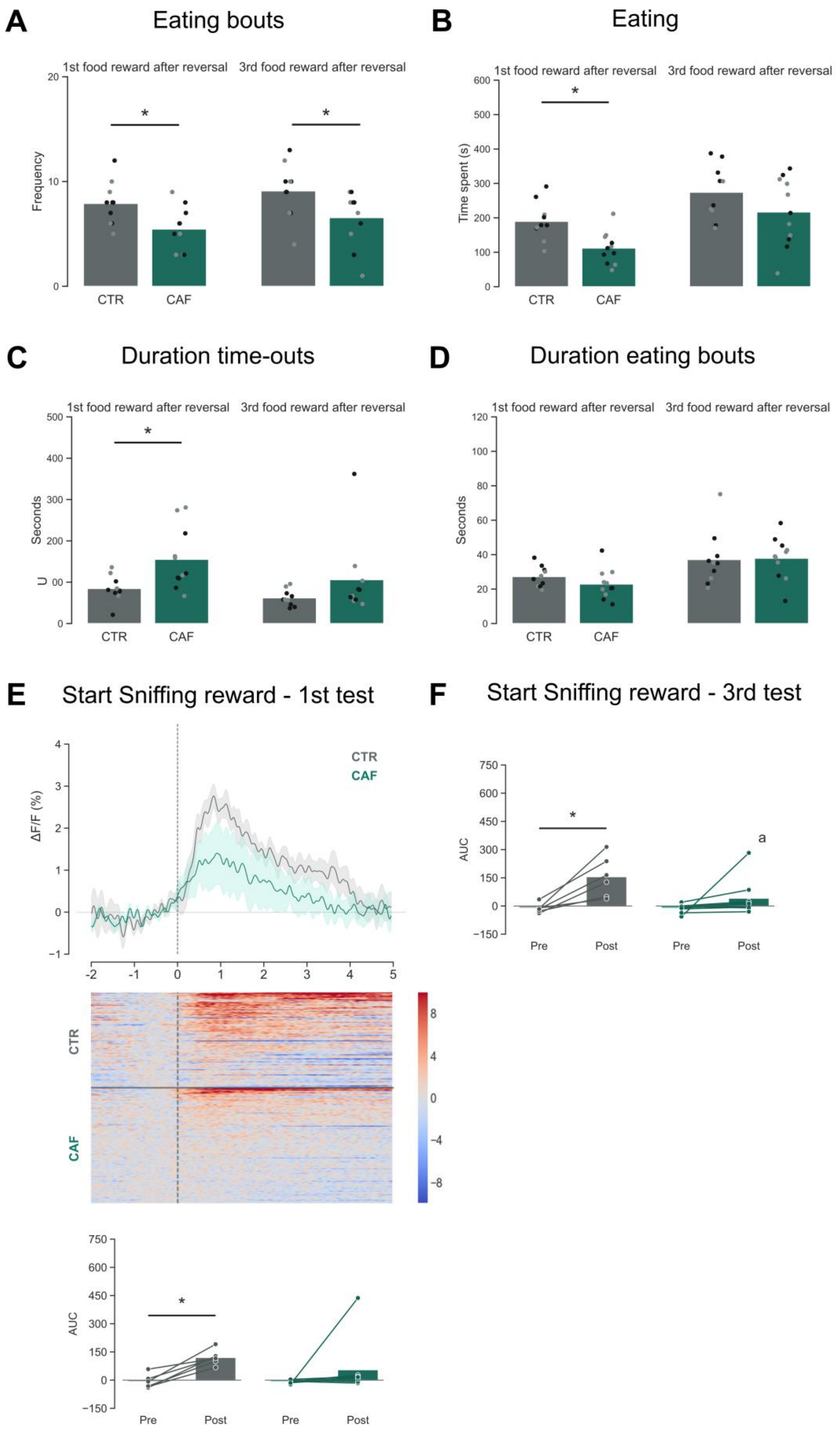
A
Eating bouts
1st food reward after reversal
3rd food reward after reversal
Frequency
CTR
CAF
B
Eating
Time spent (s)
C
Duration time-outs
Seconds
D
Duration eating bouts
E
Start Sniffing reward - 1st test
ΔF/F (%)
AUC
Pre
Post
F
Start Sniffing reward - 3rd test
a

**Figure S3.** The effects of reversal diet after Cafeteria diet exposure on the ventral tegmental area and behavioral responses during a food reward test. **A)** Time spent eating, **B)** Number of eating bouts, **C)** Mean duration of time-outs, **D)** Mean duration of eating bouts, **E)** Upper panel: ΔF/F aligned to the start of sniffing behavior. Middle panel: Heat maps of ΔF/F aligned to the start of sniffing behavior. Lower panel: AUC pre and post sniffing behavior, **F)** before (pre) and after (post) sniffing behavior. **All** Figures show rats on control (CTR, grey color bars and lines) and cafeteria (CAF, green color bars and lines) diets during the reward phase of the $1^{st}$ and/or $3^{rd}$ behavioral reward test with a food reward post-reversal diet. The dots in the behavioral data have colors representing their body weight category (grey: lighter than average, black: heavier than average). The lines in the ΔF/F figures represent the mean and shaded areas the SEM. CTR n=6, CAF n=9 for fiber photometry data and CTR n= 10, CAF n=9 for behavioral data. * $p<0.05$ compared to by line connected group. [a] $p<0.05$ compared to same bar of CTR.

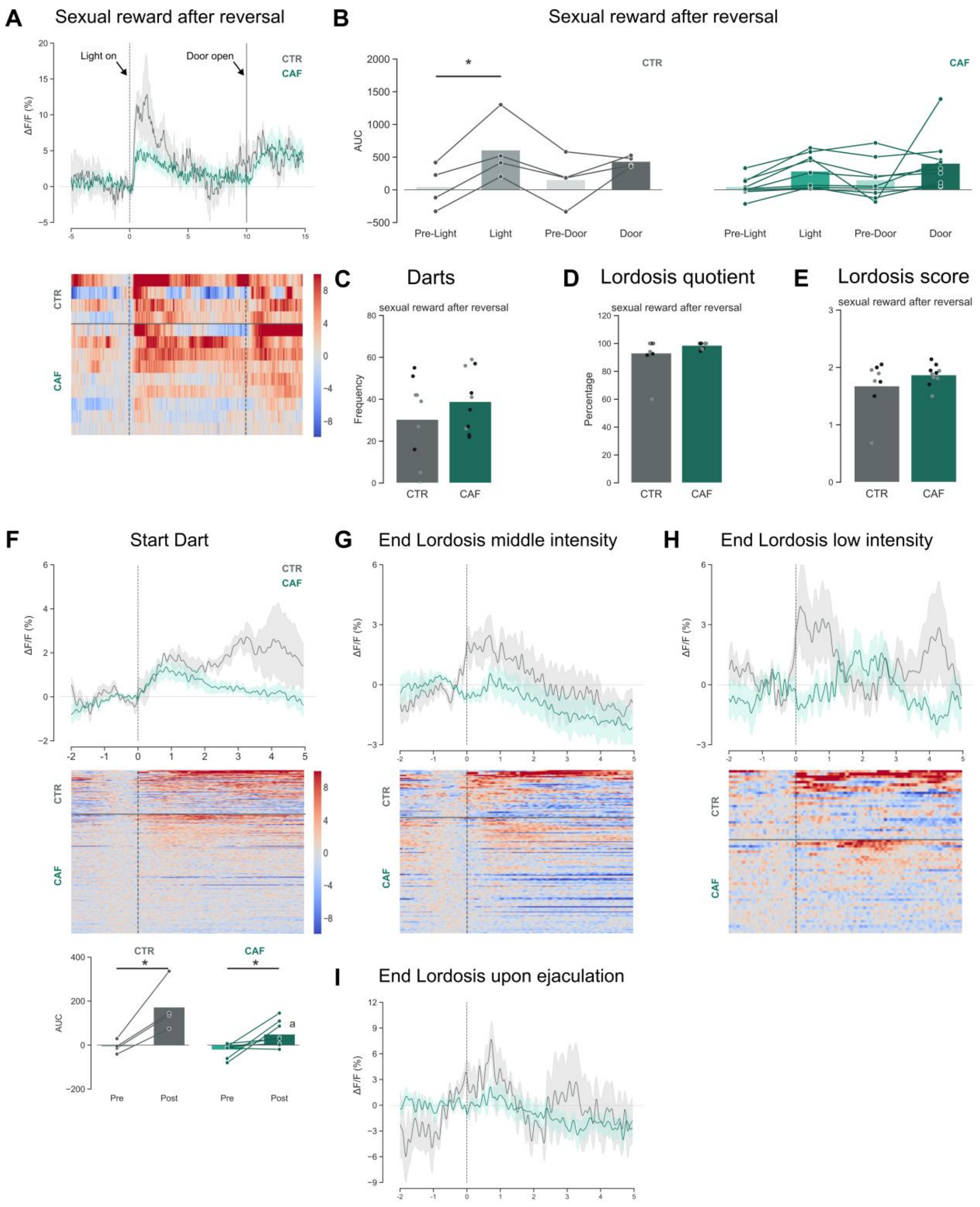


**Figure S4**. The effects of reversal diet after Cafeteria diet exposure on the ventral tegmental area and behavioral responses during a sexual reward test. A) Upper panel: ΔF/F aligned to the light cue at t=0 and door opening at t=10s. Lower panel: Heat maps of ΔF/F aligned to the light cue and door opening, B) AUC before (pre-Light) and after (Light) the light cue, and before (Pre-door) and after (Door) the door opening. C) Total number of darts, D) Lordosis quotient, E) Lordosis score, F) Upper panel: ΔF/F aligned to the start of darting behaviors. Middle panel: heat maps of ΔF/F aligned to the darting behaviors. Lower panel: AUC before (pre) and after (post) the start of darting behaviors, G) Upper panel: ΔF/F aligned to the end of middle-intensity lordosis behavior. Lower panel: heat maps of ΔF/F aligned to the end of middle-intensity lordosis behavior, H) Upper panel: ΔF/F aligned to the end of low-intensity lordosis behavior. Lower panel: heat maps of ΔF/F aligned to the end of low-intensity lordosis behavior, I) ΔF/F aligned to the end of lordosis behavior upon ejaculation. **All** Figures show rats on control (CTR, grey color bars and lines) and cafeteria (CAF, green color bars and lines) diets during the reward phase of the sexual reward test post-reversal diet. The dots in the behavioral data have colors representing their body weight category (grey: lighter than average, black: heavier than average). The lines in the ΔF/F figures represent the mean and shaded areas the SEM. CTR n=4, CAF n=9 for fiber photometry data and CTR n= 10, CAF n=9 for behavioral data. * p<0.05 compared to by line connected group. a p<0.05 compared to same bar of CTR.

### Supplementary tables

**Table S1:** Nutritional facts of the three experimental diets

| **Nutritional Facts** (per 100 g) | **Standard Chow (CTR)** (ssniff – rat/mouse – low phytoestrogen #V1554) | **High-Fat High Sugar (HE)** (RD Western Diet #D12079B) | **Cafeteria Diet (CAF)** *(4-plates together)* |
|---|---|---|---|
| **Energy density** (kcal) | 392.04 | 468.6 | 416.38 |
| **Fat (g)** | 3.4 | 21 | 22.41 |
| (saturated) | 0.54 | 13.1 | 8.38 |
| **Carbohydrates (g)** | 48.4 | 49.6 | 43.41 |
| (sugar) | 3.3 | 34.1 | 21.51 |
| **Protein (g)** | 19.1 | 19.8 | 10.0 |
| **Na (g)** | 0.23 | 0.26 | 1.59 |

**Table S2:** Nutritional values of food intake per cage in kcal and grams

| Diet | Cage | Energy (Kcal) | Fat (g) | Saturated fat (g) | Carbo-hydrate (g) | Sugar (g) | Protein (g) | Na (g) |
|---|---|---|---|---|---|---|---|---|
| CAF | 1 | 7941.59 | 395.48 | 139.76 | 769.50 | 345.12 | 228.96 | 11.32 |
| CTR | 2 | 7767.88 | 67.37 | 10.70 | 959.00 | 65.39 | 378.45 | 4.56 |
| HE | 3 | 7008.85 | 314.10 | 195.94 | 741.87 | 510.03 | 296.15 | 3.89 |
| HE | 4 | 6504.17 | 291.48 | 181.83 | 688.45 | 473.31 | 274.82 | 3.61 |
| CAF | 5 | 7815.46 | 339.07 | 119.32 | 817.82 | 333.71 | 233.02 | 8.69 |
| CTR | 6 | 7371.14 | 63.93 | 10.15 | 910.02 | 62.05 | 359.12 | 4.32 |
| CAF | 7 | 8274.65 | 325.41 | 114.59 | 930.30 | 381.45 | 254.39 | 9.15 |
| CTR | 8 | 8470.42 | 73.46 | 11.67 | 1045.73 | 71.30 | 412.67 | 4.97 |
| HE | 9 | 6909.98 | 309.67 | 193.17 | 731.40 | 502.84 | 291.97 | 3.83 |
| CAF | 10 | 9121.17 | 467.43 | 167.24 | 914.78 | 410.23 | 244.86 | 11.99 |
| HE | 11 | 7604.44 | 340.79 | 212.59 | 804.91 | 553.37 | 321.31 | 4.22 |
| CTR | 12 | 7433.47 | 64.47 | 10.24 | 917.71 | 62.57 | 362.16 | 4.36 |
| CTR | 14 | 7370.74 | 63.92 | 10.15 | 909.97 | 62.04 | 359.10 | 4.32 |
| CAF | 15 | 8469.37 | 427.03 | 155.25 | 864.16 | 437.04 | 231.68 | 11.44 |
| CAF | 16 | 8827.66 | 453.00 | 163.48 | 874.93 | 378.95 | 242.44 | 12.25 |
| HE | 17 | 6399.20 | 286.78 | 178.89 | 677.34 | 465.67 | 270.39 | 3.55 |
| CTR | 18 | 7590.29 | 65.83 | 10.45 | 937.07 | 63.89 | 369.80 | 4.45 |
| CTR | | 7584.08 ± 152.02 | 65.77 ± 1.32 | 10.45 ± 0.21 | 936.31 ± 18.77 | 63.84 ± 1.28 | 369.49 ± 7.41 | 4.45 ± 0.09 |
| CAF | | 8319.17 ± 244.03^ | 397.91 ± 25.23^* | 142.34 ± 9.53^* | 853.00 ± 27.49^ | 379.13 ± 16.77^* | 235.61 ± 4.47^* | 10.70 ± 0.65^* |
| HE | | 6779.52 ± 227.07 | 303.82 ± 10.18^* | 189.53 ± 6.35^* | 717.59 ± 24.03* | 493.35 ± 16.52^* | 286.46 ± 9.59^* | 3.76 ± 0.13 |

*Note 1:* * $p<0.05$ compared to CTR, ^ $p<0.05$ compared to HE (or CAF).

*Note 2:* Measures are based on the weight differences of the food placed and the food leftover in the cage. As always with this kind of measurements, caution should be taken because environmental factors (such a humidity or invisible food crumbs) could influence the food weight.

*Note 3:* The 1st day consumption of plate-1 food items of rat batch 1 was taken out of this analysis to match the total consumption of batch 2 who were never exposed to this extra day of food.

**Table S3:** Food consumption per cage per round for each food item.

**Biscuit consumption by CAF rats in cage 1**

| Round | Giflar kanel (g) | Brownie (g) | Marmorkake (g) | Sjokorullade (g) |
|---|---|---|---|---|
| 1 | 21.1 | 13.9 | 15.8 | 26.5 |
| 2 | 30.9 | 25.4 | 9.3 | 24 |
| 3 | 23.3 | 25.7 | 8.9 | 18.2 |
| 4 | 22 | 25.6 | 6.7 | 19.4 |
| 5 | 18.7 | 18.7 | 7.7 | 19.8 |
| 6 | 12.1 | 8.9 | 6.9 | 19.1 |
| 7 | 9.5 | 11.5 | 8.6 | 15.3 |
| 8 | 11.8 | 11.7 | 7.6 | 19.3 |
| 9 | 14.1 | 13.6 | 9 | 18.4 |
| 10 | 13.1 | 22.6 | 6.8 | 22.6 |
| 11 | 6.3 | 17.3 | 4.7 | 15.8 |
| 12 | 5.9 | 17.6 | 1.6 | 13.6 |
| 13 | 6.1 | 8.3 | 2.9 | 15 |
| **Total** | 194.9 | 220.8 | 96.5 | 247 |

**Biscuit consumption by CAF rats in cage 5**

| Round | Giflar kanel (g) | Brownie (g) | Marmorkake (g) | Sjokorullade (g) |
|---|---|---|---|---|
| 1 | 21.5 | 13 | 14.5 | 28 |
| 2 | 23.1 | 25.6 | 13.2 | 23.2 |
| 3 | 23.3 | 26.6 | 11.9 | 24.5 |
| 4 | 13.5 | 25.9 | 12.6 | 20.5 |
| 5 | 18.7 | 19 | 5.7 | 19.2 |
| 6 | 10.7 | 22.5 | 9.1 | 21.2 |
| 7 | 11.2 | 18.3 | 5.3 | 19.6 |
| 8 | 11.2 | 16.3 | 5.4 | 14.7 |
| 9 | 13.1 | 19 | 9 | 17.6 |
| 10 | 8.1 | 23.3 | 3.2 | 20.1 |
| 11 | 7.5 | 21.7 | 1.8 | 13.9 |
| 12 | 6.2 | 24 | 0.9 | 10.9 |
| 13 | 6 | 14.4 | 0.9 | 17.1 |
| **Total** | 174.1 | 269.6 | 93.5 | 250.5 |

**Biscuit consumption by CAF rats in cage 7**

| Round | Giflar kanel (g) | Brownie (g) | Marmorkake (g) | Sjokorullade (g) |
|---|---|---|---|---|
| 1 | 1.7 | 6.7 | 15.1 | 26.8 |
| 2 | 7.1 | 8.6 | 13.7 | 25.9 |
| 3 | 3.7 | 15.7 | 14.4 | 21.9 |
| 4 | 12.5 | 17.2 | 10.5 | 14.5 |
| 5 | 9.1 | 14.5 | 8.7 | 25.7 |
| 6 | 9.4 | 20.7 | 12 | 21.6 |
| 7 | 5.8 | 14.2 | 4.2 | 8.6 |
| 8 | 8.6 | 10.5 | 6 | 16 |
| 9 | 7.6 | 18.5 | 4.6 | 12.9 |
| 10 | 9.7 | 18.5 | 7 | 18.3 |
| 11 | 10.4 | 18.5 | 4.5 | 10.5 |
| 12 | 3.2 | 11.7 | 1.4 | 8.9 |
| 13 | 5.2 | 8.3 | 3.5 | 13.5 |
| **Total** | 94 | 183.6 | 105.6 | 225.1 |

**Biscuit consumption by CAF rats in cage 10**

| Round | Giflar kanel (g) | Brownie (g) | Marmorkake (g) | Sjokorullade (g) |
|---|---|---|---|---|
| 1 | - | 26.7 | 20.3 | 26.6 |
| 2 | 17.8 | 15 | 14.3 | 9.2 |
| 3 | 9.9 | 12.9 | 13.6 | 8.2 |
| 4 | 8.3 | 25.9 | 13.4 | 9.1 |
| 5 | 3.8 | 20 | 9.9 | 17.8 |
| 6 | 3.4 | 23.2 | 11.5 | 3.7 |
| 7 | 2.7 | 21 | 4.4 | 12 |
| 8 | 5.4 | 22.2 | 7.6 | 10.8 |
| 9 | 6.8 | 14.6 | 10 | 15.1 |
| 10 | 5.2 | 16.3 | 6.9 | 11.3 |
| 11 | 13 | 23.4 | 3.8 | 2.7 |
| 12 | 10.2 | 11 | 8.7 | 3.7 |
| 13 | 4.9 | 11.2 | 4.5 | 11.4 |
| **Total** | 91.4 | 232.2 | 124.4 | 130.2 |

**Biscuit consumption by CAF rats in cage 15**

| Round | Giflar kanel (g) | Brownie (g) | Marmorkake (g) | Sjokorullade (g) |
|---|---|---|---|---|
| 1 | - | 18.3 | 12.3 | 34.2 |
| 2 | 9.4 | 7.7 | 13.1 | 37.3 |
| 3 | 19 | 12.8 | 7.5 | 29.7 |
| 4 | 15.5 | 9.3 | 9.8 | 25.5 |
| 5 | 8.5 | 25.2 | 4.4 | 24 |
| 6 | 4.3 | 13.7 | 5.2 | 26.1 |
| 7 | 11.3 | 8.7 | 4 | 16.2 |
| 8 | 12.9 | 10.5 | 3.1 | 18.8 |
| 9 | 11.9 | 9.6 | 10.8 | 21 |
| 10 | 5.9 | 1.8 | 7.8 | 15.6 |
| 11 | 4.9 | 19.4 | 5.1 | 9.9 |
| 12 | 6.8 | 7.6 | 5.7 | 11.5 |
| 13 | 2.9 | 3.3 | 2.5 | 11.6 |
| **Total** | 113.3 | 147.9 | 91.3 | 281.4 |

**Biscuit consumption by CAF rats in cage 16**

| Round | Giflar kanel (g) | Brownie (g) | Marmorkake (g) | Sjokorullade (g) |
|---|---|---|---|---|
| 1 | - | 22.4 | 8.8 | 29 |
| 2 | 23.1 | 18.9 | 13.3 | 28 |
| 3 | 7.8 | 15 | 7.5 | 19.9 |
| 4 | 8.9 | 18.8 | 4.1 | 13.5 |
| 5 | 7.8 | 28.8 | 9.8 | 19.7 |
| 6 | 5.5 | 15.1 | 5.3 | 16.6 |
| 7 | 9.6 | 17.1 | 7.2 | 17.3 |
| 8 | 7.1 | 18 | 4.5 | 13.1 |
| 9 | 10.5 | 12.2 | 11.2 | 16.4 |
| 10 | 8 | 14.5 | 7.9 | 11.3 |
| 11 | 9.2 | 15.1 | 4.2 | 7.3 |
| 12 | 5.1 | 6.7 | 6.9 | 9.3 |
| 13 | 4.1 | 6.1 | 1.9 | 7.1 |
| **Total** | 106.7 | 208.7 | 90.7 | 208.5 |

**Processed meat consumption by CAF rats in cage 1**

| Round | Bacon (g) | Falukorv (g) | Salami (g) | Pølse (g) |
|---|---|---|---|---|
| 1 | 21.4 | 20.5 | 10.9 | 8.3 |
| 2 | 18.1 | 20.4 | 6.9 | 14.5 |
| 3 | 24 | 24.8 | 9.4 | 14.7 |
| 4 | 23.8 | 21.1 | 5.9 | 17.2 |
| 5 | 23.6 | 18.6 | 8.8 | 22.9 |
| 6 | 22.9 | 18.1 | 8.6 | 19.5 |
| 7 | 20.4 | 23 | 8.1 | 17.9 |
| 8 | 26.1 | 23.8 | 8 | 16.2 |
| 9 | 6.6 | 18.6 | 6.6 | 16.5 |
| 10 | 26.2 | 20.1 | 7.2 | 17.1 |
| 11 | 28.3 | 19 | 4.8 | 14.3 |
| 12 | 5.7 | 15.7 | 5.7 | 10.4 |
| 13 | 22.5 | 15.7 | 2.9 | 11 |
| **Total** | 269.6 | 259.4 | 93.8 | 200.5 |

**Processed meat consumption by CAF rats in cage 5**

| Round | Bacon (g) | Falukorv (g) | Salami (g) | Pølse (g) |
|---|---|---|---|---|
| 1 | 14.2 | 7.2 | 4.4 | 6 |
| 2 | 18.3 | 7.1 | 3 | 4.7 |
| 3 | 17.4 | 9.5 | 3.3 | 6.9 |
| 4 | 23.6 | 10.2 | 3.1 | 5.2 |
| 5 | 21.2 | 7.9 | 2.9 | 22.7 |
| 6 | 22.7 | 7.5 | 4.9 | 23.9 |
| 7 | 19.4 | 3.7 | 3.5 | 23.5 |
| 8 | 23.5 | 8.7 | 3.4 | 4.4 |
| 9 | 24.7 | 7.7 | 1.5 | 9.6 |
| 10 | 25.6 | 8.1 | 1.9 | 5.2 |
| 11 | 25.1 | 5 | 2.1 | 3 |
| 12 | 19.7 | 7.9 | 2.1 | 3 |
| 13 | 21.8 | 5.3 | 1.3 | 3.8 |
| **Total** | 277.2 | 95.8 | 37.4 | 121.9 |

**Processed meat consumption by CAF rats in cage 7**

| Round | Bacon (g) | Falukorv (g) | Salami (g) | Pølse (g) |
|---|---|---|---|---|
| 1 | 15 | 4.6 | 3.3 | 6.1 |
| 2 | 13.8 | 6.3 | 2.3 | 4.5 |
| 3 | 16.5 | 6.4 | 2.3 | 6.2 |
| 4 | 21 | 8.4 | 2.7 | 4.3 |
| 5 | 17.1 | 4.3 | 2.1 | 4.6 |
| 6 | 19.4 | 6 | 4.1 | 4.3 |
| 7 | 20.2 | 6.2 | 4.7 | 5 |
| 8 | 20.5 | 5.7 | 4.3 | 4.3 |
| 9 | 23 | 6.3 | 4.1 | 2.9 |
| 10 | 22.8 | 13.6 | 3.2 | 5.3 |
| 11 | 23.5 | 5.8 | 3.5 | 3.4 |
| 12 | 19.5 | 10.8 | 2.3 | 2.4 |
| 13 | 21.8 | 3.5 | 2.1 | 3.6 |
| **Total** | 254.1 | 87.9 | 41 | 56.9 |

**Processed meat consumption by CAF rats in cage 10**

| Round | Bacon (g) | Falukorv (g) | Salami (g) | Pølse (g) |
|---|---|---|---|---|
| 1 | - | 8.5 | 7.8 | 10.8 |
| 2 | 21.6 | 7.6 | 10.5 | 19.2 |
| 3 | 19.9 | 10.7 | 9.5 | 7.6 |
| 4 | 17.6 | 8.6 | 7.2 | 6.5 |
| 5 | 16.3 | 10.7 | 7.8 | 6.2 |
| 6 | 16.1 | 9.1 | 8.3 | 4.7 |
| 7 | 20.4 | 9.9 | 7.4 | 8.6 |
| 8 | 22.7 | 11.2 | 6 | 6.9 |
| 9 | 23.2 | 10.6 | 4.1 | 9.1 |
| 10 | 20.2 | 14.5 | 5.2 | 6.5 |
| 11 | 20.3 | 15.3 | 4.9 | 4.4 |
| 12 | 18.1 | 9.7 | 4 | 4.3 |
| 13 | 14.6 | 10 | 12.4 | 3.4 |
| **Total** | 231 | 126.4 | 95.1 | 98.2 |

**Processed meat consumption by CAF rats in cage 15**

| Round | Bacon (g) | Falukorv (g) | Salami (g) | Pølse (g) |
|---|---|---|---|---|
| 1 | - | 23.4 | 6.5 | 16.4 |
| 2 | 17.5 | 16.2 | 2.9 | 11.8 |
| 3 | 19 | 20.5 | 3.7 | 5.2 |
| 4 | 24.5 | 16.3 | 3.3 | 6.4 |
| 5 | 24.3 | 20.4 | 4.3 | 7 |
| 6 | 25.4 | 21.3 | 7.8 | 7.9 |
| 7 | 29.4 | 14.3 | 9 | 12.4 |
| 8 | 26.4 | 19.6 | 7.5 | 10.8 |
| 9 | 25.4 | 20.6 | 7 | 7.8 |
| 10 | 25.7 | 22.7 | 5.8 | 6.5 |
| 11 | 28.2 | 26.3 | 4.6 | 5 |
| 12 | 22.7 | 16.3 | 3 | 4.9 |
| 13 | 20.8 | 17.9 | 2.4 | 5 |
| **Total** | 289.3 | 255.8 | 67.8 | 107.1 |

**Processed meat consumption by CAF rats in cage 16**

| Round | Bacon (g) | Falukorv (g) | Salami (g) | Pølse (g) |
|---|---|---|---|---|
| 1 | - | 17 | 5.2 | 12 |
| 2 | 20.8 | 18.6 | 4.3 | 10.6 |
| 3 | 22.7 | 12.5 | 6 | 4.9 |
| 4 | 25.9 | 12.1 | 4.4 | 5 |
| 5 | 28.2 | 11.9 | 5 | 3.5 |
| 6 | 24.2 | 12.9 | 6.9 | 4.5 |
| 7 | 27.7 | 9.6 | 5.2 | 7.1 |
| 8 | 29.6 | 9 | 5.3 | 5.6 |
| 9 | 23.2 | 10 | 6 | 5.4 |
| 10 | 26.3 | 111.4 | 5.6 | 5.5 |
| 11 | 22.1 | 10.5 | 5.7 | 5.3 |
| 12 | 14.7 | 6.7 | 4.8 | 3.1 |
| 13 | 17.4 | 2.8 | 3.6 | 4.1 |
| **Total** | 282.8 | 245 | 68 | 76.6 |

**Candy and cookie consumption by CAF rats in cage 1**

| Round | Surt skum (g) | Sjokolade kjeks (g) | Marshmallows (g) | Oreo (g) |
|---|---|---|---|---|
| 1 | 2.3 | 4.3 | 18 | 0 |
| 2 | 0 | 0.5 | 13.6 | 1 |
| 3 | 0.1 | 0.7 | 11.7 | 0 |
| 4 | 0.6 | 1.5 | 5.7 | 0 |
| 5 | 0 | 1.1 | 8.7 | 0.6 |
| 6 | 0.3 | 0.5 | 3.2 | 2.2 |
| 7 | 0.3 | 4.9 | 7.2 | 0.4 |
| 8 | 0 | 5.6 | 7.7 | 0.8 |
| 9 | 0 | 0.9 | 8.3 | 0 |
| 10 | 0 | 3.4 | 7.8 | 1.3 |
| 11 | 1.6 | 2.6 | 2.9 | 1.3 |
| 12 | 0 | 0.5 | 2.6 | 0 |
| 13 | 0 | 2.3 | 3.3 | 0 |
| **Total** | 5.2 | 28.8 | 100.7 | 7.6 |

**Candy and cookie consumption by CAF rats in cage 5**

| Round | Surt skum (g) | Sjokolade kjeks (g) | Marshmallows (g) | Oreo (g) |
|---|---|---|---|---|
| 1 | 0 | 4.5 | 17.4 | 0 |
| 2 | 0 | 1.5 | 18 | 2.1 |
| 3 | 0.1 | 1.2 | 6.8 | 2.6 |
| 4 | 1.1 | 5.6 | 1.5 | 1.6 |
| 5 | 0.1 | 2.9 | 2.9 | 3.5 |
| 6 | 0 | 2.1 | 2.2 | 3 |
| 7 | 0 | 1.4 | 1.8 | 1.7 |
| 8 | 0 | 4.1 | 0.5 | 1.6 |
| 9 | 0 | 2.2 | 0.4 | 0 |
| 10 | 0 | 4.9 | 0 | 4.6 |
| 11 | 0.5 | 2.3 | 0.3 | 0.7 |
| 12 | 0 | 0.8 | 1 | 1.7 |
| 13 | 0 | 1.7 | 0.6 | 0.4 |
| **Total** | 1.8 | 35.2 | 53.4 | 23.5 |

**Candy and cookie consumption by CAF rats in cage 7**

| Round | Surt skum (g) | Sjokolade kjeks (g) | Marshmallows (g) | Oreo (g) |
|---|---|---|---|---|
| 1 | 0 | 1.4 | 14.9 | 0 |
| 2 | 0 | 3.8 | 16.9 | 1.6 |
| 3 | 0.5 | 2.1 | 16.2 | 1.1 |
| 4 | 1.5 | 1 | 18.3 | 5.3 |
| 5 | 5.3 | 2.6 | 16.9 | 0.5 |
| 6 | 0.1 | 1.4 | 15.7 | 0 |
| 7 | 0 | 2.3 | 18.1 | 0.2 |
| 8 | 0 | 4 | 19.5 | 1.1 |
| 9 | 0 | 2 | 16.9 | 0.2 |
| 10 | 0.3 | 4.7 | 16.9 | 2.7 |
| 11 | 0 | 4.9 | 8.9 | 0.8 |
| 12 | 0.2 | 6.6 | 9.8 | 0 |
| 13 | 0 | 6.5 | 10.6 | 0.5 |
| **Total** | 7.9 | 43.3 | 199.6 | 14 |

**Candy and cookie consumption by CAF rats in cage 10**

| Round | Surt skum (g) | Sjokolade kjeks (g) | Marshmallows (g) | Oreo (g) |
|---|---|---|---|---|
| 1 | - | 8.9 | 15.5 | 11 |
| 2 | 0.1 | 12.2 | 22.4 | 16.2 |
| 3 | 0.1 | 15 | 18.4 | 12.1 |
| 4 | 0 | 17.1 | 13.2 | 13.4 |
| 5 | 0.2 | 18.9 | 12.8 | 12.8 |
| 6 | 0 | 10.8 | 14.3 | 14.9 |
| 7 | 0 | 13.7 | 11.3 | 13.2 |
| 8 | 0 | 7.4 | 11 | 11 |
| 9 | 0 | 14 | 10.9 | 8.5 |
| 10 | 0.3 | 13.8 | 10.1 | 4.8 |
| 11 | 0.2 | 11.4 | 7.1 | 4.4 |
| 12 | 0 | 6.9 | 6.4 | 6.6 |
| 13 | 0 | 7.1 | 2.7 | 5.9 |
| **Total** | 0.9 | 157.2 | 153.4 | 134.8 |

**Candy and cookie consumption by CAF rats in cage 15**

| Round | Surt skum (g) | Sjokolade kjeks (g) | Marshmallows (g) | Oreo (g) |
|---|---|---|---|---|
| 1 | - | 6.9 | 20.1 | 10.7 |
| 2 | 0 | 10.7 | 24.9 | 3.2 |
| 3 | 0 | 10.2 | 26.9 | 1.7 |
| 4 | 0 | 9.9 | 23.8 | 2.7 |
| 5 | 0 | 15.8 | 20.5 | 4.4 |
| 6 | 0.1 | 13.8 | 21.2 | 1.1 |
| 7 | 1.6 | 8.2 | 20.4 | 0.6 |
| 8 | 0 | 9.7 | 21.1 | 2.9 |
| 9 | 0 | 7.6 | 15.8 | 0 |
| 10 | 0.1 | 11.5 | 16.8 | 1.7 |
| 11 | 0.2 | 10.4 | 12.7 | 0.2 |
| 12 | 0 | 5 | 13.9 | 0 |
| 13 | 0 | 5.7 | 8.2 | 0 |
| **Total** | 2 | 119.7 | 246.3 | 29.2 |

**Candy and cookie consumption by CAF rats in cage 16**

| Round | Surt skum (g) | Sjokolade kjeks (g) | Marshmallows (g) | Oreo (g) |
|---|---|---|---|---|
| 1 | - | 7.8 | 18.4 | 11 |
| 2 | 0 | 12.2 | 13.6 | 1.2 |
| 3 | 1.1 | 12.4 | 10.4 | 10 |
| 4 | 2.2 | 9.1 | 11.1 | 8.5 |
| 5 | 1.3 | 15.4 | 15.2 | 10.9 |
| 6 | 1.5 | 6.4 | 7.3 | 4.6 |
| 7 | 0.1 | 8.4 | 6.9 | 5.4 |
| 8 | 0.8 | 9.1 | 6.3 | 5.6 |
| 9 | 0 | 6.6 | 7.1 | 8.4 |
| 10 | 0 | 8.7 | 9.4 | 6.5 |
| 11 | 0.1 | 11.9 | 7.1 | 5.8 |
| 12 | 0 | 5.2 | 6.1 | 4.1 |
| 13 | 0 | 1.4 | 2.9 | 6.5 |
| **Total** | 7.1 | 114.6 | 121.8 | 88.5 |

**Salty snack consumption by CAF rats in cage 1**

| Round | Cheez doodles (g) | Lu Tuc (g) | Goldfish (g) | Bacon svor (g) |
|---|---|---|---|---|
| 1 | 0.6 | 5.5 | 3.9 | 0.4 |
| 2 | 0.1 | 2.1 | 5.2 | 0 |
| 3 | 0.2 | 2 | 2.2 | 0 |
| 4 | 0.7 | 0.9 | 0 | 0 |
| 5 | 2.9 | 1.6 | 1.8 | 0 |
| 6 | 3.4 | 0.6 | 1.4 | 0.9 |
| 7 | 7.9 | 0.3 | 1.5 | 0 |
| 8 | 3.3 | 0.5 | 1.4 | 1.3 |
| 9 | 5 | 0.5 | 3.2 | 2.4 |
| 10 | 4.1 | 2.3 | 3.1 | 2.3 |
| 11 | 4 | 0.2 | 2 | 0.4 |
| 12 | 1.9 | 0.9 | 3.9 | 0.4 |
| 13 | 1.9 | 0.8 | 2 | 1.4 |
| **Total** | 36 | 18.2 | 31.6 | 9.5 |

**Salty snack consumption by CAF rats in cage 5**

| Round | Cheez doodles (g) | Lu Tuc (g) | Goldfish (g) | Bacon svor (g) |
|---|---|---|---|---|
| 1 | 0 | 2.9 | 0 | 0.2 |
| 2 | 0.4 | 1.1 | 0 | 0 |
| 3 | 0.2 | 1 | 0 | 0 |
| 4 | 0.5 | 0.5 | 0.7 | 0.4 |
| 5 | 0.6 | 2 | 0 | 0 |
| 6 | 0.6 | 2 | 0 | 0 |
| 7 | 0.6 | 0 | 1.4 | 0.9 |
| 8 | 0 | 1.1 | 1.4 | 3.3 |
| 9 | 1 | 1 | 1.6 | 2.8 |
| 10 | 5.8 | 1.9 | 0.2 | 2.8 |
| 11 | 4.6 | 1 | 0 | 1.5 |
| 12 | 1.2 | 3.2 | 0 | 0 |
| 13 | 1.3 | 0.8 | 0 | 1.8 |
| **Total** | 16.8 | 18.5 | 5.3 | 13.7 |

**Salty snack consumption by CAF rats in cage 7**

| Round | Cheez doodles (g) | Lu Tuc (g) | Goldfish (g) | Bacon svor (g) |
|---|---|---|---|---|
| 1 | 0 | 0.2 | 4.2 | 3.4 |
| 2 | 1.2 | 6.3 | 7.7 | 5 |
| 3 | 5.3 | 7.3 | 9.5 | 0.6 |
| 4 | 5.2 | 5.2 | 5.3 | 2.6 |
| 5 | 5.3 | 3.2 | 4.4 | 0 |
| 6 | 1.9 | 3.9 | 4.6 | 1.5 |
| 7 | 3.1 | 4.4 | 3 | 2.6 |
| 8 | 1.9 | 2.1 | 3.9 | 6.2 |
| 9 | 0.5 | 5.5 | 2.4 | 6.3 |
| 10 | 7.6 | 3.5 | 1.9 | 6.4 |
| 11 | 6 | 1.5 | 1.1 | 0.8 |
| 12 | 3.6 | 2.9 | 4.2 | 2.6 |
| 13 | 5.6 | 1 | 0.5 | 3.4 |
| **Total** | 47.2 | 47 | 52.7 | 38 |

**Salty snack consumption by CAF rats in cage 10**

| Round | Cheez doodles (g) | Lu Tuc (g) | Goldfish (g) | Bacon svor (g) |
|---|---|---|---|---|
| 1 | - | 3.6 | 7.8 | 6.5 |
| 2 | 14.2 | 13.2 | 4.4 | 5.6 |
| 3 | 17.9 | 11.8 | 1.8 | 8.7 |
| 4 | 20.2 | 17.1 | 0 | 10.7 |
| 5 | 18.8 | 9.6 | 0.7 | 10.4 |
| 6 | 14.8 | 4.1 | 0 | 10.1 |
| 7 | 14 | 6.5 | 0 | 8.8 |
| 8 | 11.3 | 6.4 | 0 | 5.4 |
| 9 | 5.1 | 3 | 0.9 | 9.9 |
| 10 | 13.4 | 1.5 | 1.4 | 6.7 |
| 11 | 9 | 5 | 0.3 | 4.4 |
| 12 | 4.4 | 2 | 7.8 | 5.5 |
| 13 | 8.2 | 2.7 | 0.1 | 3.8 |
| **Total** | 151.3 | 86.5 | 25.2 | 96.5 |

**Salty snack consumption by CAF rats in cage 15**

| Round | Cheez doodles (g) | Lu Tuc (g) | Goldfish (g) | Bacon svor (g) |
|---|---|---|---|---|
| 1 | - | 7.8 | 7.8 | 0.4 |
| 2 | 9.5 | 15.8 | 12.8 | 0 |
| 3 | 5 | 11.3 | 4.5 | 3.9 |
| 4 | 6.5 | 9.9 | 0 | 4.8 |
| 5 | 4.1 | 7.7 | 0 | 5.2 |
| 6 | 5.4 | 0.8 | 0.4 | 7.1 |
| 7 | 5.8 | 5 | 0 | 8.8 |
| 8 | 5.7 | 4.4 | 0.6 | 5.2 |
| 9 | 5.1 | 1.5 | 0 | 7.7 |
| 10 | 8.6 | 0.2 | 0.5 | 7.6 |
| 11 | 10.8 | 9.4 | 0 | 4.4 |
| 12 | 4.9 | 1.9 | 0.5 | 4.6 |
| 13 | 6.5 | 0.3 | 0 | 7 |
| **Total** | 77.9 | 76 | 27.1 | 66.7 |

**Salty snack consumption by CAF rats in cage 16**

| Round | Cheez doodles (g) | Lu Tuc (g) | Goldfish (g) | Bacon svor (g) |
|---|---|---|---|---|
| 1 | - | 7.9 | 7.9 | 1.7 |
| 2 | 12.1 | 10.9 | 3.9 | 0.3 |
| 3 | 16.1 | 9.1 | 5.9 | 5.8 |
| 4 | 11.9 | 9.3 | 4.2 | 6.7 |
| 5 | 14.7 | 9.2 | 1.1 | 5.6 |
| 6 | 13.9 | 7.6 | 2.7 | 7.9 |
| 7 | 13.5 | 6.4 | 2.8 | 7.9 |
| 8 | 11 | 6.3 | 3.1 | 6.5 |
| 9 | 11.3 | 5.4 | 2.3 | 13.3 |
| 10 | 7.9 | 4.9 | 2.3 | 8.6 |
| 11 | 14 | 7.2 | 4.4 | 2.7 |
| 12 | 6 | 5.5 | 0.7 | 2.2 |
| 13 | 5.8 | 5.2 | 0.9 | 3.6 |
| **Total** | 138.2 | 89.7 | 41.3 | 69.2 |

**Standard chow consumption by CAF rats in cage 1**

| Round | Plate-1 (g) | Plate-2 (g) | Plate-3 (g) | Plate-4 (g) |
|---|---|---|---|---|
| 1 | 13.4 | 10.3 | 1.9 | 3.6 |
| 2 | 3.1 | 8.2 | 14.7 | 0.1 |
| 3 | 0.2 | 5.2 | 1.3 | 0 |
| 4 | 5.4 | 17.5 | 14 | 0 |
| 5 | 5.8 | 11.9 | 7.6 | 10.7 |
| 6 | 12 | 13.3 | 8.6 | 8.7 |
| 7 | 3.9 | 13.7 | 10.1 | 9.2 |
| 8 | 3.7 | 12.2 | 5.2 | 1.6 |
| 9 | 2.2 | 12.7 | 15.8 | 3 |
| 10 | 5.7 | 2.4 | 11.3 | 1.1 |
| 11 | 6 | 12 | 8.3 | 8.5 |
| 12 | 10.5 | 12.1 | 5.5 | 3.6 |
| 13 | 4.9 | 9.5 | 10.2 | 6.5 |
| **Total** | 76.8 | 141 | 114.5 | 56.6 |

**Standard chow consumption by CAF rats in cage 5**

| Round | Plate-1 (g) | Plate-2 (g) | Plate-3 (g) | Plate-4 (g) |
|---|---|---|---|---|
| 1 | 21.4 | 17.2 | 8.5 | 12.9 |
| 2 | 7.5 | 5.2 | 9.7 | 10.1 |
| 3 | 2.3 | 14.5 | 10.6 | 9.8 |
| 4 | 7.4 | 17.2 | 7.4 | 10.5 |
| 5 | 6 | 9 | 15.5 | 16.4 |
| 6 | 8.9 | 11.5 | 17 | 13.7 |
| 7 | 4.1 | 16.7 | 12.1 | 14.4 |
| 8 | 6 | 15 | 20.7 | 8.9 |
| 9 | 9.4 | 12.3 | 19.4 | 7.6 |
| 10 | 4.2 | 13 | 18.2 | 12.4 |
| 11 | 5.1 | 15.5 | 9.1 | 9.4 |
| 12 | 8.7 | 9.2 | 9.9 | 8.3 |
| 13 | 4.4 | 15.4 | 12.9 | 11.3 |
| **Total** | 95.4 | 171.7 | 171 | 145.7 |

**Standard chow consumption by CAF rats in cage 7**

| Round | Plate-1 (g) | Plate-2 (g) | Plate-3 (g) | Plate-4 (g) |
|---|---|---|---|---|
| 1 | 22.3 | 28.3 | 21.1 | 10.2 |
| 2 | 24.4 | 21.6 | 4.8 | 12.1 |
| 3 | 19.7 | 23.2 | 5.7 | 13.7 |
| 4 | 14.9 | 23.2 | 4.2 | 10.1 |
| 5 | 14.1 | 16.5 | 1.8 | 8.1 |
| 6 | 14.4 | 15.2 | 4.4 | 11.5 |
| 7 | 15.3 | 14.2 | 11.8 | 14.5 |
| 8 | 14.4 | 15.4 | 13.2 | 6.1 |
| 9 | 7.4 | 11.6 | 15.7 | 13.9 |
| 10 | 4.8 | 11.7 | 12.5 | 7.5 |
| 11 | 3.4 | 10.3 | 9.7 | 11 |
| 12 | 9.7 | 6.3 | 10 | 9.4 |
| 13 | 8.1 | 15.4 | 10.7 | 10.7 |
| **Total** | 172.9 | 212.9 | 125.6 | 138.8 |

**Standard chow consumption by CAF rats in cage 10**

| Round | Plate-1 (g) | Plate-2 (g) | Plate-3 (g) | Plate-4 (g) |
|---|---|---|---|---|
| 1 | - | 6.4 | 3.1 | 4.7 |
| 2 | 0.2 | 5.8 | 9.2 | 6.2 |
| 3 | 4.1 | 23.5 | 4.6 | 0.1 |
| 4 | 6.6 | 1.9 | 7.6 | 2.3 |
| 5 | 4.3 | 5 | 1.9 | 0 |
| 6 | 13.4 | 7.4 | 12 | 2.8 |
| 7 | 8.9 | 0.5 | 6.8 | 3 |
| 8 | 3.8 | 5.2 | 9.7 | 3.1 |
| 9 | 1.3 | 5.9 | 8.6 | 2.5 |
| 10 | 2.9 | 2.8 | 9.3 | 4.3 |
| 11 | 5.9 | 6.9 | 9.6 | 1 |
| 12 | 7.4 | 4.9 | 7.6 | 4.6 |
| 13 | 6.7 | 14 | 10.6 | 7 |
| **Total** | 65.5 | 90.2 | 100.6 | 41.6 |

**Standard chow consumption by CAF rats in cage 15**

| Round | Plate-1 (g) | Plate-2 (g) | Plate-3 (g) | Plate-4 (g) |
|---|---|---|---|---|
| 1 | - | 5.8 | 5.4 | 7.4 |
| 2 | 0.2 | 14.8 | 9.6 | 8.7 |
| 3 | 4.9 | 1.1 | 6.3 | 3.9 |
| 4 | 1 | 0.9 | 6.7 | 5.6 |
| 5 | 6.3 | 2.2 | 9.9 | 5.5 |
| 6 | 4.7 | 5.3 | 4.6 | 7.3 |
| 7 | 7.8 | 0.7 | 2.5 | 7 |
| 8 | 5.6 | 7 | 3 | 2.6 |
| 9 | 2.6 | 5.9 | 6.4 | 3 |
| 10 | 1.6 | 4.5 | 1.9 | 0 |
| 11 | 7.5 | 4 | 5.6 | 1.8 |
| 12 | 3.9 | 5.3 | 6.7 | 1.3 |
| 13 | 2.4 | 5.8 | 7.2 | 6.3 |
| **Total** | 48.5 | 63.3 | 75.8 | 60.4 |

**Standard chow consumption by CAF rats in cage 16**

| Round | Plate-1 (g) | Plate-2 (g) | Plate-3 (g) | Plate-4 (g) |
|---|---|---|---|---|
| 1 | - | 10.9 | 8.3 | 10 |
| 2 | 0.3 | 0 | 11.1 | 9.7 |
| 3 | 0.8 | 7.6 | 9.8 | 5.6 |
| 4 | 0 | 4.6 | 10.2 | 6.5 |
| 5 | 0 | 9.9 | 11.5 | 6.3 |
| 6 | 0 | 7.8 | 7.3 | 6.2 |
| 7 | 0 | 0 | 9.2 | 8.5 |
| 8 | 2.3 | 4.6 | 8 | 4.9 |
| 9 | 1.7 | 1.7 | 12.3 | 4.9 |
| 10 | 2.4 | 8.3 | 13.5 | 2.1 |
| 11 | 1.3 | 5.6 | 9.9 | 3.9 |
| 12 | 1.8 | 5.3 | 13.3 | 3.3 |
| 13 | 1.7 | 7.8 | 13.1 | 7.6 |
| **Total** | 12.3 | 74.1 | 137.5 | 79.5 |

**Standard chow consumption by CTR rats in cage 2**

| Round | Plate-1 (g) | Plate-2 (g) | Plate-3 (g) | Plate-4 (g) |
|---|---|---|---|---|
| 1 | 42.1 | 36.4 | 36.9 | 40.8 |
| 2 | 26.9 | 37.5 | 33.5 | 34.7 |
| 3 | 35.2 | 38.9 | 31.1 | 38.2 |
| 4 | 36.2 | 34.7 | 40.9 | 37.7 |
| 5 | 37 | 27.3 | 39.1 | 40.1 |
| 6 | 34 | 34.9 | 32.5 | 34.9 |
| 7 | 42.2 | 35.8 | 43.6 | 36.3 |
| 8 | 41 | 34.6 | 36.2 | 41.9 |
| 9 | 36.5 | 38 | 36.8 | 32.8 |
| 10 | 33.7 | 37.7 | 36.3 | 42.4 |
| 11 | 36.2 | 34.3 | 31.4 | 26.6 |
| 12 | 27 | 35 | 22.5 | 18.9 |
| 13 | 27.1 | 28.7 | 26.2 | 34.3 |
| **Total** | 455.1 | 453.8 | 447 | 459.6 |

**Standard chow consumption by CTR rats in cage 6**

| Round | Plate-1 (g) | Plate-2 (g) | Plate-3 (g) | Plate-4 (g) |
|---|---|---|---|---|
| 1 | 38.3 | 34.2 | 30.4 | 38.7 |
| 2 | 29.3 | 31.2 | 32.8 | 35.5 |
| 3 | 37.4 | 37.5 | 26.2 | 36.4 |
| 4 | 41 | 35.8 | 34.1 | 30.5 |
| 5 | 44.3 | 31.9 | 37.7 | 39.1 |
| 6 | 36.5 | 34.4 | 33.1 | 33.2 |
| 7 | 43.2 | 36.3 | 36 | 36 |
| 8 | 38.2 | 35.5 | 35 | 33.3 |
| 9 | 40.2 | 39 | 35.8 | 34.5 |
| 10 | 38.8 | 33.1 | 36.5 | 57.4 |
| 11 | 38.8 | 32.8 | 29.2 | 30.3 |
| 12 | 27.6 | 28.3 | 23.6 | 25.5 |
| 13 | 34.3 | 35.7 | 16.1 | 19.9 |
| **Total** | 487.9 | 445.7 | 406.5 | 450.3 |

**Standard chow consumption by CTR rats in cage 8**

| Round | Plate-1 (g) | Plate-2 (g) | Plate-3 (g) | Plate-4 (g) |
|---|---|---|---|---|
| 1 | 47 | 36 | 39.3 | 37.2 |
| 2 | 38.6 | 44.5 | 36.6 | 40.4 |
| 3 | 43.9 | 49.5 | 37 | 41.9 |
| 4 | 47.6 | 35.3 | 44.7 | 29.7 |
| 5 | 54.5 | 34.3 | 43.2 | 42.1 |
| 6 | 43.4 | 38 | 38.1 | 33.2 |
| 7 | 47.9 | 42.8 | 40.3 | 35.8 |
| 8 | 38.5 | 38.8 | 36.4 | 38.6 |
| 9 | 42 | 41 | 38 | 36.2 |
| 10 | 45.1 | 36.5 | 39.9 | 37.5 |
| 11 | 46.6 | 36.1 | 27.4 | 31.6 |
| 12 | 32.9 | 38.2 | 15.1 | 29.4 |
| 13 | 35.3 | 36.3 | 33.3 | 39.6 |
| **Total** | 563.3 | 507.3 | 469.3 | 473.2 |

**Standard chow consumption by CTR rats in cage 12**

| Round | Plate-1 (g) | Plate-2 (g) | Plate-3 (g) | Plate-4 (g) |
|---|---|---|---|---|
| 1 | - | 37.9 | 41.7 | 31.2 |
| 2 | 24.5 | 49 | 43.2 | 34.6 |
| 3 | 36.4 | 38.7 | 30.3 | 34.6 |
| 4 | 35 | 35.3 | 44.7 | 32.1 |
| 5 | 43.5 | 38.5 | 41.6 | 33.4 |
| 6 | 44.6 | 39.7 | 38.4 | 30.5 |
| 7 | 40.6 | 33.9 | 40 | 38.3 |
| 8 | 39.9 | 34.7 | 39.2 | 30.7 |
| 9 | 27.9 | 35.2 | 34 | 32.9 |
| 10 | 34.7 | 36.5 | 32.7 | 33.2 |
| 11 | 34.4 | 41.7 | 30.1 | 22.1 |
| 12 | 23.9 | 27.6 | 27.2 | 19.5 |
| 13 | 21.9 | 27.8 | 29.4 | 25.7 |
| **Total** | 407.3 | 476.5 | 472.5 | 398.8 |

**Standard chow consumption by CTR rats in cage 14**

| Round | Plate-1 (g) | Plate-2 (g) | Plate-3 (g) | Plate-4 (g) |
|---|---|---|---|---|
| 1 | - | 35.7 | 35.1 | 28.3 |
| 2 | 41.1 | 36.9 | 36.4 | 32 |
| 3 | 36.3 | 38.7 | 35.3 | 33.2 |
| 4 | 39.4 | 38.3 | 37.9 | 29.7 |
| 5 | 37.3 | 30 | 31.4 | 40.4 |
| 6 | 35.6 | 33.6 | 35.7 | 36.8 |
| 7 | 39.1 | 38.4 | 36.9 | 39.9 |
| 8 | 32.9 | 33.1 | 29.2 | 32.9 |
| 9 | 31 | 40.2 | 31.1 | 32.5 |
| 10 | 30.6 | 35 | 30.4 | 38 |
| 11 | 31.3 | 42.1 | 25.4 | 28.7 |
| 12 | 32.3 | 31.4 | 25.4 | 25.8 |
| 13 | 28 | 28.8 | 24.6 | 31.5 |
| **Total** | 414.9 | 462.2 | 414.8 | 429.7 |

**Standard chow consumption by CAF rats in cage 18**

| Round | Plate-1 (g) | Plate-2 (g) | Plate-3 (g) | Plate-4 (g) |
|---|---|---|---|---|
| 1 | - | 34.7 | 35.9 | 32 |
| 2 | 42.4 | 39.8 | 37.8 | 39 |
| 3 | 39.1 | 38 | 34 | 44.8 |
| 4 | 34.6 | 35.1 | 40.1 | 40.1 |
| 5 | 35.2 | 34.9 | 40.3 | 34.6 |
| 6 | 32.4 | 47.2 | 38.7 | 36.2 |
| 7 | 36.9 | 35.6 | 37.3 | 39 |
| 8 | 38.7 | 39.8 | 34.8 | 40.2 |
| 9 | 32.5 | 39.4 | 39.5 | 33.7 |
| 10 | 30.4 | 36.2 | 38.1 | 37.4 |
| 11 | 38.3 | 36.6 | 36.1 | 22 |
| 12 | 26 | 29 | 30.3 | 17 |
| 13 | 22.7 | 31 | 28.2 | 27.7 |
| **Total** | 409.2 | 477.3 | 471.1 | 443.7 |

**HE pellet consumption by HE rats in cage 3**

| Round | Plate-1 (g) | Plate-2 (g) | Plate-3 (g) | Plate-4 (g) |
|---|---|---|---|---|
| 1 | 37.4 | 28.4 | 37.8 | 20.3 |
| 2 | 31.6 | 21.3 | 26.3 | 27.1 |
| 3 | 30.4 | 25.6 | 25.4 | 20.5 |
| 4 | 31.9 | 30 | 22.6 | 28.8 |
| 5 | 28.9 | 26.8 | 29 | 20.1 |
| 6 | 26.3 | 22.8 | 31.1 | 24.8 |
| 7 | 27.1 | 23.7 | 26 | 27.8 |
| 8 | 27.7 | 24.2 | 29 | 28.5 |
| 9 | 22.7 | 21.1 | 31.3 | 22.2 |
| 10 | 34.6 | 20.7 | 29 | 26.9 |
| 11 | 30.1 | 25.6 | 11.8 | 22.9 |
| 12 | 35.5 | 24.9 | 23 | 21.9 |
| 13 | 32.4 | 19.2 | 22.5 | 25.8 |
| **Total** | 396.6 | 314.3 | 344.8 | 317.6 |

**HE pellet consumption by HE rats in cage 4**

| Round | Plate-1 (g) | Plate-2 (g) | Plate-3 (g) | Plate-4 (g) |
|---|---|---|---|---|
| 1 | 36.8 | 29.4 | 28.9 | 25.5 |
| 2 | 25.9 | 27.1 | 29.5 | 22.7 |
| 3 | 24.4 | 32.1 | 25 | 27.4 |
| 4 | 25.4 | 32.2 | 29.6 | 19.6 |
| 5 | 27.2 | 23.3 | 23.9 | 27 |
| 6 | 25.9 | 21.7 | 24.5 | 27.2 |
| 7 | 25.5 | 25.8 | 27.6 | 25.1 |
| 8 | 24.1 | 22.6 | 24 | 28.2 |
| 9 | 21.4 | 26.3 | 28.3 | 26.4 |
| 10 | 23.1 | 22.7 | 25.7 | 24.6 |
| 11 | 24.3 | 21.4 | 17.6 | 21.9 |
| 12 | 24 | 23.2 | 15.3 | 17.3 |
| 13 | 24.9 | 15.1 | 24.5 | 17.8 |
| **Total** | 332.9 | 322.9 | 324.4 | 310.7 |

**HE pellet consumption by HE rats in cage 9**

| Round | Plate-1 (g) | Plate-2 (g) | Plate-3 (g) | Plate-4 (g) |
|---|---|---|---|---|
| 1 | 38.7 | 31.8 | 38.4 | 28.1 |
| 2 | 27.8 | 33.6 | 29.8 | 26 |
| 3 | 27.3 | 27 | 27.7 | 22.9 |
| 4 | 28.7 | 24.4 | 24.6 | 32.1 |
| 5 | 25.7 | 24 | 27.1 | 28.9 |
| 6 | 26.3 | 24.1 | 25.1 | 18.3 |
| 7 | 28.7 | 32.5 | 25 | 29.5 |
| 8 | 22 | 30.2 | 20.9 | 23.8 |
| 9 | 28.8 | 18.9 | 31.8 | 23.1 |
| 10 | 24.9 | 22.2 | 25.6 | 22.4 |
| 11 | 22.6 | 26 | 19.2 | 20 |
| 12 | 23.3 | 22.5 | 20.4 | 21.1 |
| 13 | 22.9 | 20.6 | 23.5 | 18.3 |
| **Total** | 347.7 | 337.8 | 339.1 | 314.5 |

**HE pellet consumption by HE rats in cage 11**

| Round | Plate-1 (g) | Plate-2 (g) | Plate-3 (g) | Plate-4 (g) |
|---|---|---|---|---|
| 1 | - | 58 | 50.3 | 44 |
| 2 | 46.8 | 32.7 | 39.7 | 30.9 |
| 3 | 33 | 30.6 | 32.1 | 32.1 |
| 4 | 28.7 | 27.6 | 37 | 23.1 |
| 5 | 27.5 | 26.4 | 30.8 | 25.6 |
| 6 | 23.9 | 29.8 | 34.3 | 36.1 |
| 7 | 27.3 | 26.9 | 28.5 | 29.5 |
| 8 | 28.6 | 30.7 | 26.4 | 29 |
| 9 | 24 | 23.9 | 26.1 | 32.3 |
| 10 | 25.4 | 26.2 | 18.6 | 29 |
| 11 | 30.8 | 24.3 | 19.8 | 17.6 |
| 12 | 22.4 | 32.7 | 20.6 | 18.7 |
| 13 | 20.8 | 25.3 | 18.6 | 22.3 |
| **Total** | 31.6 | 35.5 | 31.9 | 36.5 |

**HE pellet consumption by HE rats in cage 17**

| Round | Plate-1 (g) | Plate-2 (g) | Plate-3 (g) | Plate-4 (g) |
|---|---|---|---|---|
| 1 | - | 47.6 | 41.7 | 39 |
| 2 | 31.6 | 28.3 | 35 | 31.4 |
| 3 | 17.1 | 29.6 | 27.5 | 29.5 |
| 4 | 31.6 | 16.8 | 27.4 | 26.4 |
| 5 | 23.1 | 27.2 | 23.6 | 22 |
| 6 | 19.5 | 29.8 | 31.8 | 24.7 |
| 7 | 21.7 | 20 | 18.9 | 19.3 |
| 8 | 25.9 | 26 | 19.3 | 28.5 |
| 9 | 20.4 | 27.6 | 21.6 | 20.5 |
| 10 | 22.4 | 23.6 | 23.4 | 21 |
| 11 | 27.2 | 24.8 | 32.9 | 17.2 |
| 12 | 17.4 | 23.2 | 23.9 | 12.2 |
| 13 | 17.2 | 16.1 | 22 | 21.2 |
| **Total** | 275.1 | 340.6 | 349 | 312.9 |

*Note:* Rats in batch 2 started their diet exposure with plate 2 and therefore received 1 day of food less. This is visualized with the symbol –
